# Molecularly-Defined Hippocampal Inputs Regulate Population Dynamics in the Prelimbic Cortex to Suppress Context Fear Memory Recall

**DOI:** 10.1101/802967

**Authors:** Henry L. Hallock, Henry M. Quillian, Kristen R. Maynard, Yishan Mai, Huei-Ying Chen, Gregory R. Hamersky, Joo Heon Shin, Brady J. Maher, Andrew E. Jaffe, Keri Martinowich

## Abstract

Associating fearful events with the context in which they occur is critical for survival. Dysregulation of context-fear memory processing is a hallmark symptom of several neuropsychiatric disorders, including generalized anxiety disorder (GAD) and post-traumatic stress disorder (PTSD). Both the hippocampus and prelimbic subregion (PrL) of the medial prefrontal cortex (mPFC) have been linked with context fear memory recall in rodents, but the mechanisms by which hippocampal-prelimbic circuitry regulates this process remains poorly understood. Spatial and genetic targeting of this circuit in mice allowed us to use molecular profiling to show that hippocampal neurons with projections to the PrL (vHC-PrL projectors) are a transcriptomically-distinct sub-population that is enriched for expression of genes associated with both GAD and PTSD. We further show that stimulation of this population of vHC-PrL projectors suppresses context fear memory recall and impairs the ability of PrL neurons to dynamically distinguish between distinct phases of fear learning. Using transgenic and circuit-specific molecular targeting approaches, we demonstrate that unique patterns of activity-dependent gene transcription within vHC-PrL projectors causally regulate excitatory/inhibitory balance in the PrL during context fear memory recall. Together, our data illuminate the molecular mechanisms by which hippocampal-prelimbic circuitry regulates the retrieval of contextually-mediated fear memories.

## Introduction

Aberrant prefrontal cortex (PFC) function is implicated in many neuropsychiatric disorders, including generalized anxiety disorder (GAD)^1^, and post-traumatic stress disorder (PTSD)^2^. Patients with these disorders frequently have deficits in cognitive/behavioral domains that are supported by the PFC. For example, patients with PTSD show altered regulation of fear^3^ and deficits in emotional memory^4^. Theories of emotional memory dysregulation in PTSD have centered on deficient context processing^5, 6^, suggesting that patients with PTSD have core functional deficits in PFC circuits that are critical for relating emotional memories with the context in which they occur. Elucidating how these circuits encode context during emotional memory formation and retrieval is therefore important for understanding how changes in neural function drive fear dysregulation in these disorders.

In rodents, the prelimbic (PrL) subdivision of the medial PFC (mPFC) critically contributes to fear memory expression during recall^7^, while the infralimbic (IL) subdivision critically contributes to fear extinction^8^. Recent evidence, however, suggests that the relationship between PrL function and fear behavior is more nuanced. For example, stimulating excitatory PrL-IL connections suppresses freezing during fear recall^9^, and bulk calcium transients in the PrL predict decreases in freezing during extinction training^10^, suggesting that the PrL can flexibly encode distinct fear states during memory formation, retrieval, and extinction. The PrL is necessary for context fear memory recall^11^, raising the possibility that the PrL dynamically represents shifts in context to guide appropriate behavior across distinct phases of fear learning. The PrL additionally receives input from many brain areas, including those known to be involved in context encoding. One such brain area is the hippocampus (HC), which sends efferent projections to the PrL via neurons in the ventral CA1 subfield^12, 13^. The ventral HC (vHC) is also critical for context-dependent fear expression^14^, suggesting that PrL-projecting neurons in the vHC (vHC-PrL projectors) may be important for mediating the contextual component of emotional memories. Can PrL neurons code for contextually-dependent shifts in fear state during memory formation, recall, and extinction? If so, how do inputs from the vHC affect this process at the circuit, cellular, and molecular levels?

To answer these questions, we used molecular-genetic and systems level techniques to demonstrate that activation of monosynaptic ventral hippocampal (vHC) inputs to the PrL suppresses freezing during context fear memory recall by rendering PrL population activity less contextually-specific. Using translating ribosome affinity purification (TRAP) and RNA-sequencing, we show that vHC-PrL projection neurons express gene sets that are implicated in PTSD, and that activation of these neurons induces unique patterns of activity-dependent gene transcription associated with brain-derived neurotrophic factor (BDNF) signaling in the mPFC. Finally, we find that selective expression of BDNF in vHC-PrL projection neurons is sufficient to rescue behavioral and molecular phenotypes associated with impaired HC-PFC function and attenuate exaggerated context fear memory expression in mice.

## Results

### Molecular characterization of vHC-PrL projection neurons

Given the discrete anatomical definition of the hippocampal-prelimbic circuit, we asked whether vHC neurons with direct projections to the PrL (vHC-PrL projectors) can also be differentiated on the basis of their molecular profile. We used translating ribosome affinity purification (TRAP) to isolate HA-tagged ribosome-mRNA complexes and conduct downstream RNA-seq from vHC-PrL projectors (Fig. 1a). Specifically, we injected a Cre-expressing retrograde virus (AAVrg-hSyn1-eBFP-Cre) into the PrL of adult RiboTag mice, which allowed for Cre-mediated HA-tagging of ribosomes with bound mRNAs in monosynaptic inputs to the PrL. To confirm Cre-mediated recombination in our model, we performed anti-HA immunohistochemistry in a subset of RiboTag mice, which revealed dense labeling restricted to the CA1 subfield of the vHC in the projector group, but widespread labeling in all vHC subfields of the control group (all Syn1-expressing vHC neurons) (Fig. 1b). We next dissected out the vHC, immunoprecipitated (IP) ribosomes from HA-expressing neurons, and isolated RNA for sequencing. We used qPCR to confirm that virally-derived *Cre* and *Bfp* were selectively expressed in IP samples from the projector group (Supp. Fig. 1a), further demonstrating projector-specific expression of the RiboTag allele. RNA-seq revealed that, compared to the control group, 644 genes were enriched in projectors, and 699 genes were depleted in projectors (Fig. 1c; Supp. Table 1), and gene ontology (GO) analysis showed that genes associated with learning, memory, synaptic plasticity, and glutamatergic signaling were enriched in vHC-PrL projectors (Supp. Fig. 1a; Supp. Table 2). Projectors were also highly enriched for genes involved in post-synaptic modulation, but depleted for genes involved in pre-synaptic signaling. These results suggest that the distinct molecular composition of vHC-PrL projectors may render them poised to mediate learning and memory, possibly via direct modulation of plasticity in post-synaptic projection targets. Interestingly, we also found differential expression of several genes related to stress signaling in projectors, including enrichment of *Nr3c1*, which encodes the glucocorticoid receptor (GR) in mice, and depletion of the corticotrophin releasing factor binding protein gene *Crhbp*. Given that learning, memory, and stress-associated genes were enriched in mouse projectors, we next asked whether these genes are also implicated in disorders that feature dysregulation of fear expression in humans, specifically PTSD and GAD. Using the Harmonizome database, we found significant enrichment of our differentially-expressed genes among genes previously associated with both GAD (e.g., *Far2*; log2FC = 0.57, adjusted *p* = 0.019, *Plk2;* log2FC = 0.69, adjusted *p* = 6.2e^-6) and PTSD (e.g., *Jun;* log2FC = 0.75, adjusted *p* = 8.02e^-5), supporting a role for molecular regulation of hippocampal-prefrontal function in these disorders (adjusted *p* for GAD overlap = 2.5^-5; adjusted *p* for PTSD overlap = 0.004; all genes listed in Supp. Table 3).

**Figure 1.**
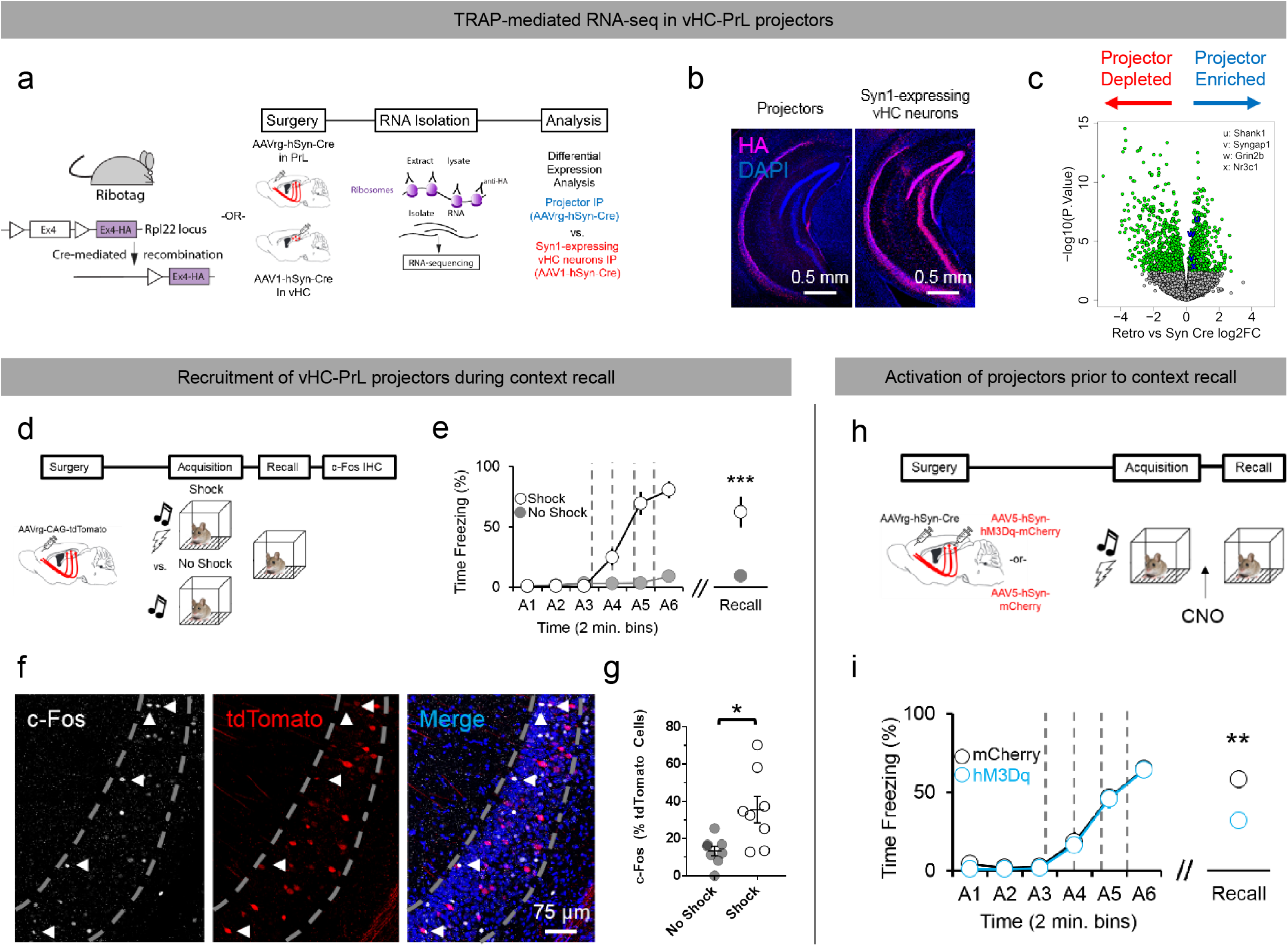
Activation of molecularly-distinct vHC-PrL projectors suppresses context fear memory recall. *(a)* Schematic of timeline and experimental design for RNA-sequencing of projectors vs. all Syn1-expressing vHC neurons. *(b)* HA protein expression in the vHC of Rpl22^HA^ animals injected with AAVrg-hSyn1-emBFP-Cre in the PrL (left panel), and AAV1-hSyn1-emBFP-Cre in the vHC (right panel). *(c)* Volcano plot showing significantly enriched genes in projectors (red dots, right of 0 on x-axis) and significantly enriched genes in Syn1-expressing vHC neurons (red dots, left of 0 on x-axis). *Shank1* log2(FC) = 0.65, adjusted *p* = 0.0000187, *Syngap1* log2FC = 0.31, adjusted *p* = 0.007, *Grin2b* log2FC = 0.3, adjusted *p* = 0.0002, *Nr3c1* log2FC = 0.45, adjusted *p* = 0.024. *(d)* Schematic of timeline and experimental design for identification of projectors recruited during context fear recall. *(e)* Mice that received a shock during conditioning froze at significantly higher levels than mice that did not receive a shock during both conditioning (*F*(5,30) = 84.26, *p* < 0.0001, mixed-factorial ANOVA) and context recall (*t*(6) = 7.075, *p* = 0.0004, unpaired t-test). *(f)* Example confocal z-projections from ventral CA1 showing tdTomato and c-Fos co-expression. *(g)* A significantly larger proportion of projectors (tdTomato+ neurons) in ventral CA1 co-express context recall-induced c-Fos in shocked mice vs. non-shocked mice (*t*(14) = 2.933, *p* = 0.01, unpaired t-test; n mice = 4 per group, n images = 2 per mouse). *(h)* Schematic of timeline and experimental design for excitation of vHC-PrL projectors prior to context fear recall. *(i)* Synthetic excitation of projectors prior to context recall significantly reduces freezing relative to viral (mCherry) controls (*t*(26) = 2.917, *p* = 0.007, unpaired t-test; n mice = 20/mCherry, 8/hM3Dq).

### Stimulation of vHC-PrL projection neurons suppresses expression of context fear memories

To elaborate a functional role for vHC-PrL projectors, we quantified their recruitment during context fear memory recall. Specifically, we injected a retrograde virus encoding a fluorescent reporter (AAVrg-CAG-tdTomato) into the PrL of adult wild-type (w/t) mice to label monosynaptic inputs, and then performed c-Fos immunohistochemistry to identify tdTomato+ neurons in ventral CA1 that were activated during context fear recall in mice that were fear conditioned (shock group), versus controls (no shock group) (Fig. 1d). As expected, mice in the shock group froze at significantly higher levels during both conditioning and context recall (Fig. 1e). The proportion of tdTomato+ neurons that co-expressed c-Fos (Fig. 1f) was significantly higher in the shock group, as compared to the no shock group (Fig. 1g), even as total number of c-Fos+ neurons did not significantly differ between groups (Supp. Fig. 1b). To causally link function in vHC-PrL projectors with context fear recall, we injected a retrograde virus expressing Cre-recombinase (AAVrg-hSyn1-emBFP-Cre) into the PrL, along with a virus encoding for either Cre-dependent expression of the excitatory DREADD receptor hM3Dq (AAV5-hSyn1-DIO-hM3Dq-mCherry) or the mCherry reporter alone (AAV5-hSyn1-DIO-mCherry) into the vHC of w/t mice, allowing us to selectively stimulate vHC-PrL projectors prior to context recall. Mice in both hM3Dq and mCherry groups were fear conditioned and administered clozapine-N-oxide (CNO; 5 mg/kg, i.p.) 45 minutes prior to context recall to engage the hM3Dq receptor (Fig. 1h). mCherry expression in the vHC of both groups was mainly restricted to the CA1 subfield, with more limited labeling in the ventral subiculum (Supp. Fig. 1c). Synthetic activation of vHC-PrL projectors significantly decreased freezing during context recall (Fig. 1i), suggesting that vHC-PrL projectors regulate the suppression of context fear memory expression.

### Stimulation of vHC-PrL projectors inhibits flexible encoding of distinct fear memory stages in PrL neurons

To understand how PrL neurons encode behavior across different stages of fear learning, we compared calcium dynamics within individual neurons in PrL across distinct fear memory phases. To do this we injected AAV1-hSyn1-GCaMP6f into the PrL and imaged calcium activity from 20-100 PrL neurons simultaneously across conditioning, context fear recall, and extinction training through an endoscopic lens with a miniature head-mounted microscope (57 ± 12.65 neurons per animal, mean ± s.e.m.; Fig. 2a). Importantly, we were able to reliably register imaged neurons included in our analyses across all fear memory phases, allowing us to directly compare freezing-related activity in the same neuron as a function of conditioning (Fig. 2b). The majority of PrL neurons imaged were active across fear memory phases (Fig. 2c), suggesting that the PrL dynamically represents behavior during fear learning, recall, and extinction. We found that many PrL neurons dynamically represented freezing bouts, showing either increased or decreased calcium activity during freezing onset (Fig. 2d for examples of a freezing-activated and freezing-inhibited neuron). For each fear memory phase, populations of PrL neurons exhibited a heterogeneous pattern of freezing-related activity; however, all imaged PrL neurons changed their activity patterns during freezing bouts as a function of fear memory phase (Fig. 2e), indicating that dynamic shifts in PrL population activity differentially represent fear state during immobility.

**Figure 2.**
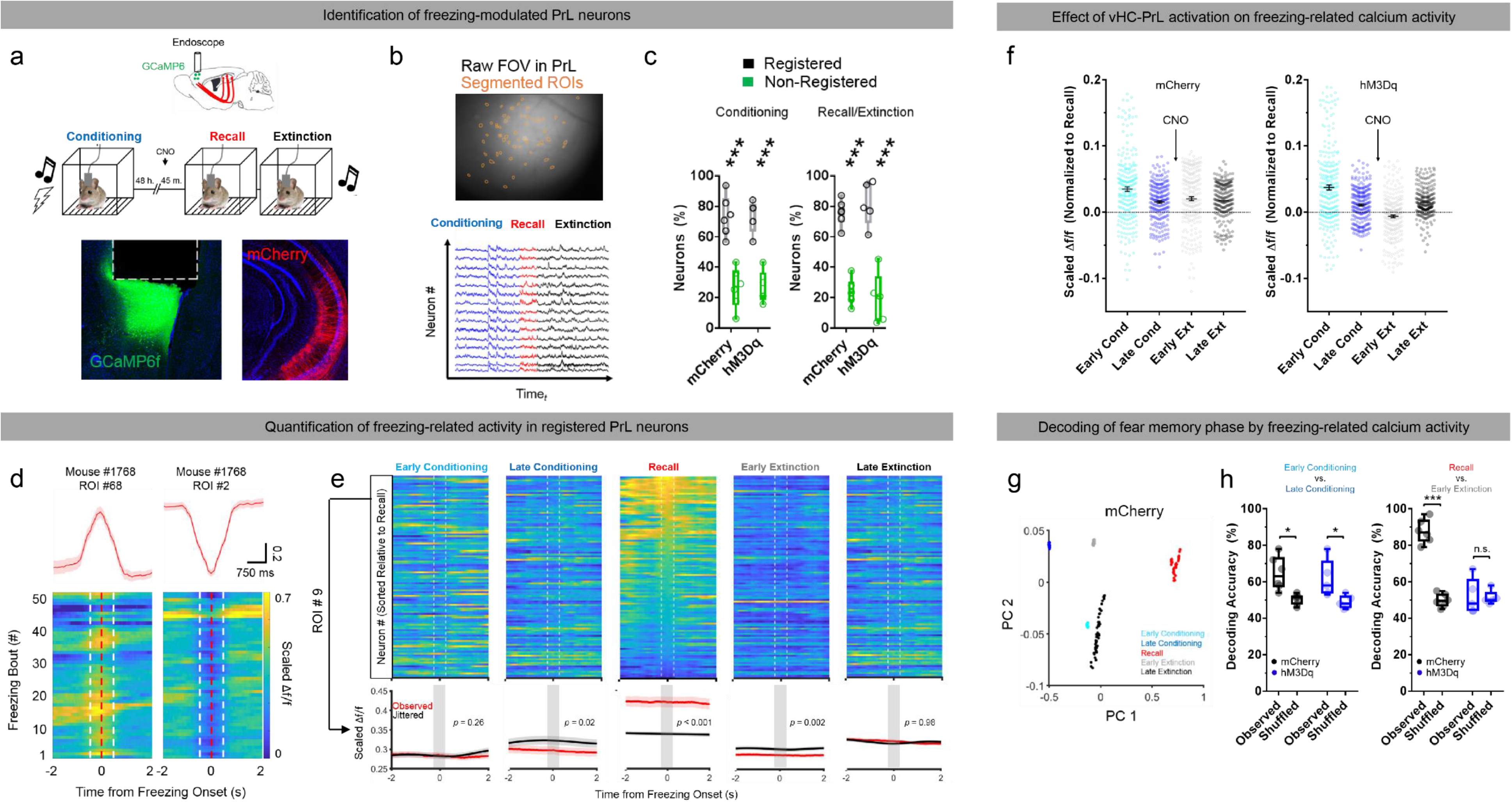
vHC-PrL projectors alter population dynamics in the PrL during context recall and extinction training. *(a)* Schematic of experimental design. (*Top*) The excitatory DREADD receptor hM3Dq was expressed in vHC-PrL projectors, and GCaMP6f was expressed in the PrL for simultaneous imaging of PrL neurons during fear conditioning, context recall, and extinction training with a miniature one-photon microscope. (*Bottom*) Example confocal z-projection from the PrL showing GCaMP6f expression (green) and an endoscopic lens track (dashed line), and the vHC showing mCherry expression in projectors. *(b) (Top)* Example field of view from the PrL showing raw GCaMP6f fluorescence, with individual ROIs (single neurons) superimposed in orange. *(Bottom)* Example traces extracted from the PrL of one mouse registered across conditioning, recall, and extinction training. *(c)* Example of a PrL neuron that shows increased activity (“freezing-increased”; *left panel*), and a PrL neuron that shows decreased activity (“freezing-decreased; *right panel*) during freezing bouts *(d)* A significant proportion of PrL neurons were able to be registered across fear phases in both mCherry and hM3Dq mice during both conditioning ((*F*(1,18) = 82.64, *p* < 0.0001, main effect of registration category) and recall/extinction training (*F*(1,18) = 120.4, *p* < 0.0001, main effect of registration category for 2 (condition) x 2 (registration category) ANOVA) *(e) (Top row)* Averaged activity from 86 simultaneously imaged PrL neurons from one mouse across early conditioning (pre-shock), late conditioning (post-shock), context recall, early extinction, and late extinction during freezing bouts. Neurons are sorted relative to their Δf/f value during context recall freezing onset (dashed white lines). *(Bottom row)* Observed and jittered (peri-freezing) activity traces from one neuron in the population. This neuron varies its freezing-related activity as a function of fear epoch, reflecting the tendency of the population to shift its freezing-related activity patterns across fear learning, recall, and extinction. *(f)* On average, calcium activity during freezing bouts is higher during early conditioning, late conditioning, early extinction, and late extinction compared to recall (black dashed line at 0 on y-axis) in mCherry control mice (left panel). In hM3Dq mice, CNO injections caused freezing-related activity in PrL neurons to be more similar between early extinction and recall (closer to black dashed line), an effect that was not observed during other fear memory phases (right panel; *F*(4) = 15.69, *p* = 1.2^e-12, significant group x fear memory phase interaction for linear mixed effects model). *(g)* Principal components analysis (PCA) on population calcium activity during freezing reveals robust separation of clusters along the first two principal component axes as a function of fear memory phase in mCherry mice (each dot on graph represents calcium activity of all registered neurons in one time bin during freezing bouts). *(h)* Decoding accuracy of a linear classifier using freezing-related calcium activity to predict fear memory phase (observed) is significantly above chance (shuffled data) when distinguishing between early and late conditioning for both groups (left panel; mCherry *p* = 0.027, hM3Dq *p* = 0.049), but falls to chance levels when distinguishing between context recall and early extinction in the hM3Dq group (right panel; mCherry *p* < 0.0001, hM3Dq *p* = 0.995, Sidak’s multiple comparison tests).

Given that stimulation of vHC-PrL projectors decreases freezing during fear recall, we hypothesized that projector stimulation might impact behavior by altering PrL representations during freezing. To investigate this question, we injected a combination of viruses to induce expression of either the excitatory hM3Dq receptor or an mCherry control selectively in vHC-PrL projectors, and administered CNO prior to context recall while simultaneously imaging calcium dynamics in the PrL during fear behavior. Unilateral activation of vHC-PrL projectors did not alter the number of freezing bouts during any fear memory phase (Supp Fig. 2a), but did impact patterns of freezing-related activity in ipsilateral PrL neurons. Specifically, we found that proportions of neurons belonging to each of the three categories differed between mCherry and hM3Dq groups during recall and extinction, but not during conditioning (Supp. Fig. 2b). To quantify variability of PrL neurons across fear memory phases at the population level, we used a linear mixed effects model with an interaction term between fear memory phase and group, and mouse and neuron as random terms. We found a significant fear memory phase-by-group interaction that was largely driven by group differences in freezing-related calcium activity during context recall and early extinction training. Specifically, our model showed that mean freezing-related calcium activity increased from context recall to early extinction training in mCherry controls, and that this effect was blunted in hM3Dq animals (Fig. 2f), indicating that projector activation renders the PrL less able to flexibly represent distinct fear memory phases during freezing.

To further determine whether PrL populations distinguish between fear memory phases during freezing, we used freezing-related calcium activity in all imaged neurons from mice in the mCherry group across fear memory phases as input for principal components analysis (PCA). We found that PrL ensemble activity during freezing tightly separated into distinct clusters, and that these clusters were highly separated along the first two principal component axes as a function of fear memory phase (Fig. 2g). To quantify whether differences in freezing-related PrL activity are sufficient to predict fear memory phase, we trained a support vector machine (SVM) classifier to distinguish between early and late conditioning, or context recall and early extinction training freezing bouts based on PrL population calcium activity. We found that decoding accuracy of the classifier was higher than chance (shuffled data) in both groups when distinguishing between early and late conditioning, but fell to chance levels in hM3Dq animals when tasked with choosing between context recall and early extinction (Fig. 2h). Taken together, our data suggest that individual PrL neurons flexibly encode the fear state of the animal independently of overt behavior, and that vHC inputs to the PrL suppress context fear recall by rendering PrL population states more rigid between recall and extinction training.

### vHC-PrL projector activation induces distinct activity-dependent gene expression profiles in the mPFC

Because input from the vHC impacts patterns of neural activity in the PrL at the cellular level, we next aimed to identify how efferent input from the vHC regulates neural activity at the molecular level in the PrL. To investigate this question, we expressed hM3Dq or an mCherry control selectively in vHC-PrL projectors and administered CNO to assess effects of projector activation on gene expression in the mPFC (Fig. 3a). We used qPCR to confirm that activation of vHC-PrL projectors induced immediate early gene (IEG) expression in bulk mPFC tissue, and found significant enrichment (∼10-30 fold increases) of several IEGs (*Arc*, *Fos*, and *Npas4*) in the mPFC of the CNO+hM3Dq group, as compared to both control groups (Fig. 3b).

**Figure 3.**
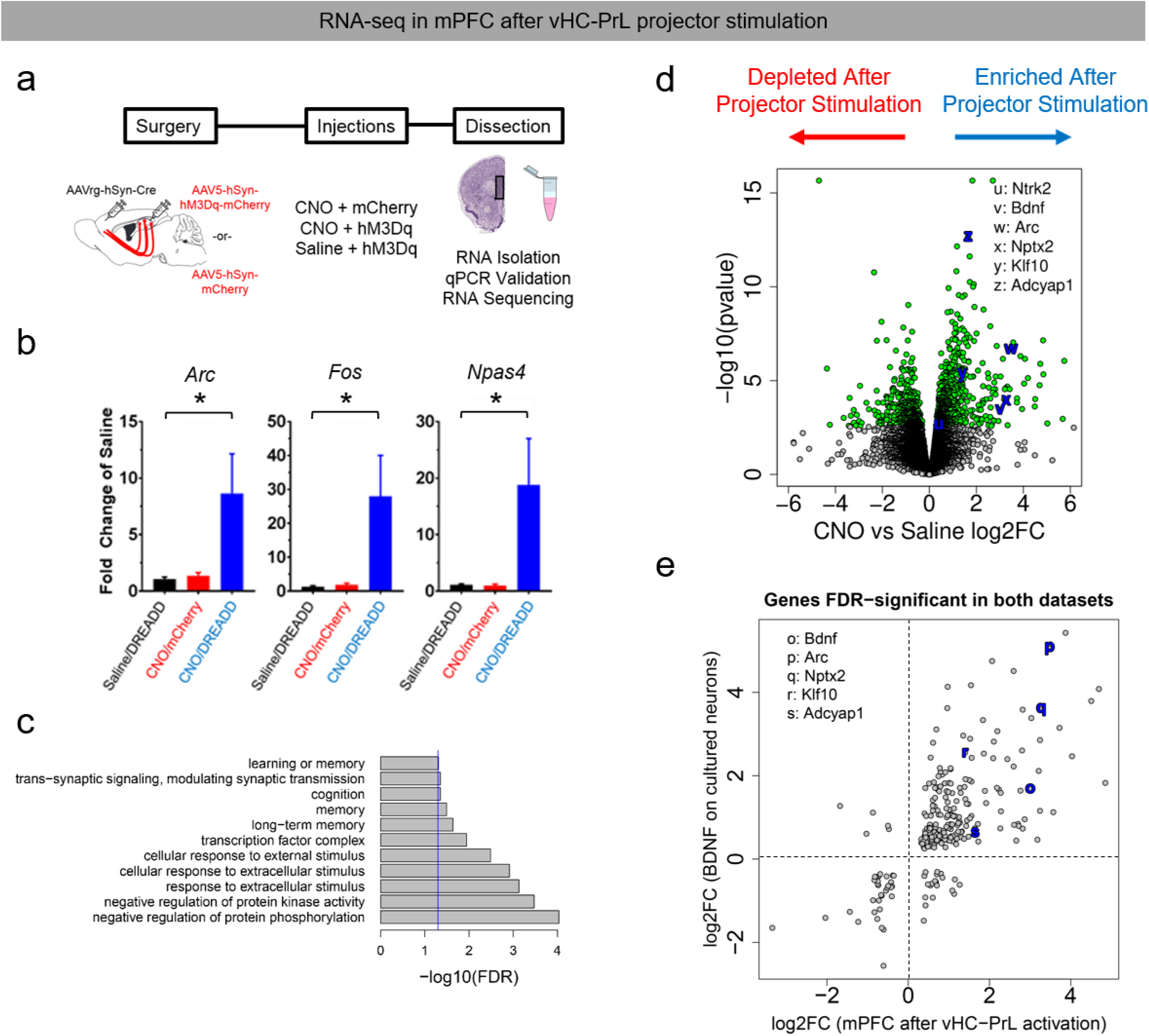
Stimulation of vHC-PrL projectors alters activity-dependent gene expression in the PrL. *(a)* Schematic of timeline and experimental design. *(b)* qPCR validates that immediate early genes (IEGs) are up-regulated in the PrL following stimulation of projectors, but not after CNO injections alone (*F*(2,11) = 3.991, *p* = 0.0497 for *Arc*, *F*(2,11) = 4.213, *p* = 0.0438 for *Fos*, *F*(2,11) = 4.112, *p* = 0.0464 for *Npas4*, one-way ANOVAs). *(c)* GO enrichment analysis reveals significant up-regulation of genes that code for proteins necessary for learning, memory, cognition, long-term memory, and synaptic modulation following activation of projectors. *(d)* Volcano plot showing significantly differentially-expressed down-regulated (left of 0 on x-axis) and up-regulated (right of 0 on x-axis) genes (red dots; FDR < 0.1) in the PrL following projector activation. Genes related to BDNF-TrkB signaling are labeled with blue letters. *Ntrk2* log2FC = 0.4, adjusted *p* = 0.044, *Bdnf* log2FC = 0.84, adjusted *p* = 0.014, *Arc* log2FC = 0.67, adjusted *p* = 7.21e^-5, *Nptx2* log2FC = 0.85, adjusted *p* = 0.0068, *Klf10* log2FC = 0.31, adjusted *p* = 0.0007, *Adcyap1* log2FC = 1.65, adjusted *p* = 1.08e^-9. *(e)* Scatter plot showing directional overlap between differentially-expressed genes in our data set and differentially-expressed genes in a published data set from cultured neurons following BDNF application (*p* < 0.0001; *Χ*^2^ test).

To identify gene sets in the mPFC that are significantly regulated by vHC inputs, we performed RNA-seq on isolated mPFC RNA from CNO+hM3Dq and saline+hM3Dq groups. Of the 20,463 expressed genes in our data set, we identified 1,156 genes that were differentially expressed between the two groups at FDR < 1% (Fig. 3c; Supp. Table 4). Of these differentially expressed genes, 914 were significantly down-regulated in the CNO+hM3Dq group, while the remaining 242 were significantly up-regulated. We confirmed that IEG upregulation measured with qPCR repeated by RNA-seq in the CNO+hM3Dq group (adjusted *p* < 0.001 for *Arc*, *Fos*, and *Npas4*). We next performed GO enrichment analysis on the subset of 242 genes that were significantly up-regulated in the CNO+hM3Dq group, and found that vHC-PrL stimulation induced mRNAs for genes that encode proteins critically involved in learning, memory, and cognition (Fig. 3d; Supp. Table 5). We also observed that this enriched gene set encoded proteins involved in modulation of synaptic transmission, transcription factor binding, and cellular responses to extracellular stimuli, supporting the hypothesis that vHC-PrL projections influence learned fear expression by modifying plasticity in post-synaptic targets. Given that synaptic transmission and transcription factor binding was significantly influenced by activation of projectors, we hypothesized that molecules synthesized in projectors that affected PrL function would be critically involved in modulation of synaptic plasticity. On further examination of our enriched gene set, we noticed that several genes directly involved in the brain-derived neurotrophic factor (BDNF)-TrkB signaling pathway were significantly up-regulated (Fig. 3e), including *Bdnf* itself, as well as *Ntrk2*, which encodes BDNF’s cognate receptor, TrkB. This result indicates that projector activation affects BDNF-TrkB signaling in the PrL during context fear memory recall, which is in line with previous literature demonstrating a critical role for BDNF signaling in hippocampal-prefrontal circuitry and fear behavior^15, 16^. To more conclusively link BDNF function with differentially expressed genes in our data set, we compared fold changes in significantly regulated genes from our data with a publicly available data set of significantly regulated genes following application of BDNF to cultured cortical neurons^17^. We found highly significant directional overlap of fold changes in differentially expressed genes between the two data sets (Fig. 3f), including *Arc* and *Ntrk2*, further implicating BDNF-TrkB signaling as a key regulator of function in the vHC-PrL circuit.

### Context fear recall induces distinct molecular phenotypes in the PrL of mutant mice with decreased activity-dependent BDNF signaling

To relate BDNF-TrkB signaling with PrL function during fear behavior, we examined cell type-specific patterns of activation following context fear recall in mutant mice with disruption of activity-dependent BDNF production (-e4 mice). In these mice, knock-in of an eGFP-STOP cassette to the *Bdnf* exon IV locus causes disruption of BDNF production from promoter IV, while leaving production from other *Bdnf* promoters intact^18^ (Fig. 4a). We fear conditioned w/t and –e4 mice, and re-exposed them to the conditioning chamber for assessment of context fear memory recall 48 h later (Fig. 4b). No significant difference in freezing was observed between genotypes during conditioning, but –e4 mice froze at significantly higher levels than w/t controls during context recall (Fig. 4c). Ninety minutes following context recall, we sacrificed mice and extracted brains for analysis of fear-related patterns of gene expression in the PrL and IL. We used single-molecule *in situ* hybridization to identify excitatory neurons (*Slc17a7* probe) recruited during context fear recall (*Fos* probe) that could be directly modulated by BDNF (*Ntrk2* probe) (Fig. 4d). We found that a higher proportion of excitatory cells receiving BDNF (*Slc17a7*+/*Ntrk2*+) in the PrL co-expressed *Fos* in -e4 mice, indicating that this population of excitatory PrL neurons is recruited to a greater degree during context recall in mutant animals. Conversely, we observed that the same population of *Slc17a7*+/*Ntrk2*+ neurons in the IL, which is necessary for fear extinction, co-expressed *Fos* to a significantly lesser degree in -e4 animals following context fear recall (Fig. 4e).

**Figure 4.**
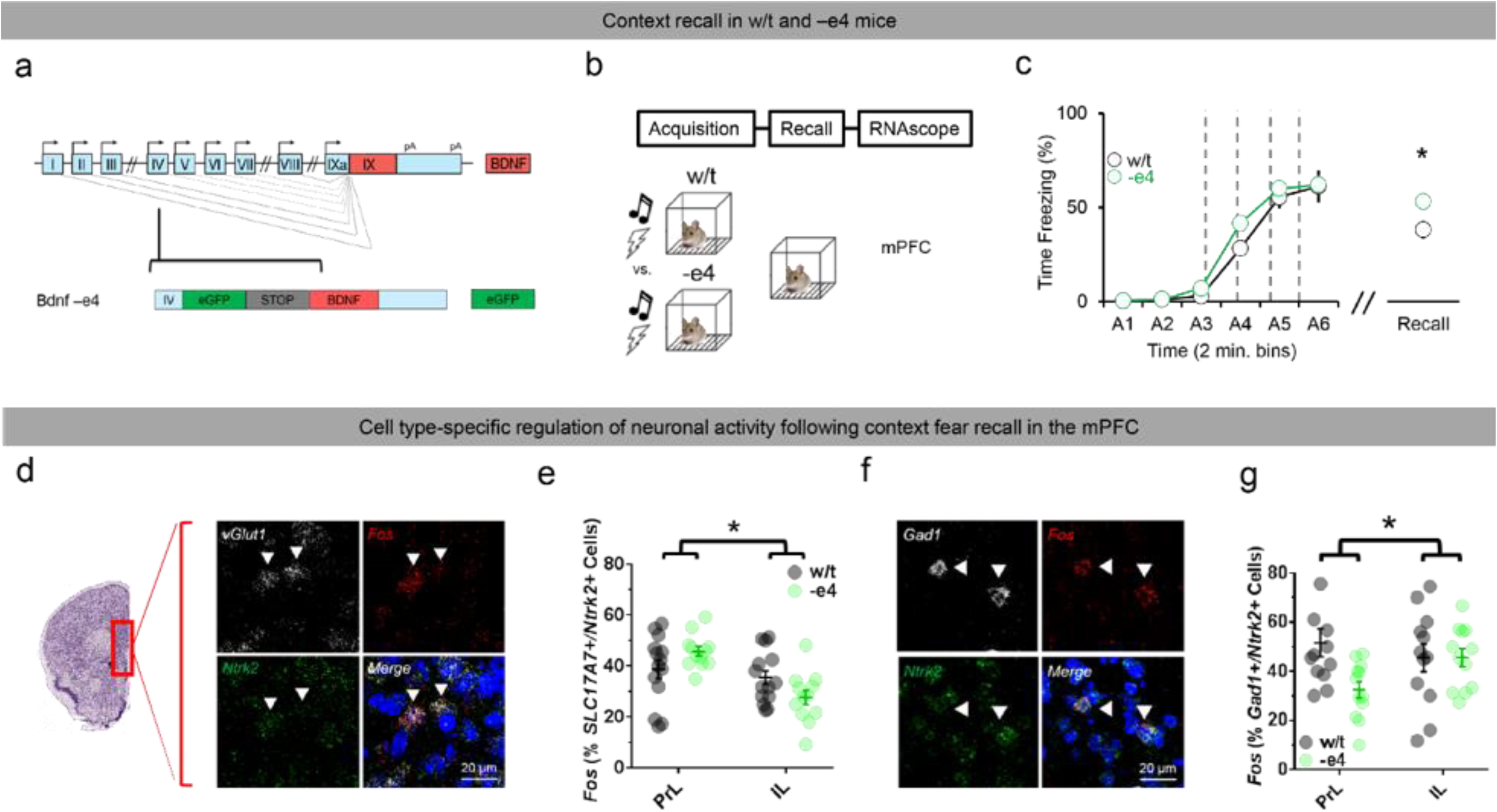
Altered context fear recall and cell type-specific gene expression in the mPFC of mice with decreased activity-dependent BDNF production. *(a)* Schematic illustrating insertion of eGFP-STOP cassette downstream of promoter IV of the *Bdnf* gene in mice. This mutation leads to mice with decreased production of BDNF from promoter IV-derived transcripts (-e4 mice). *(b)* Schematic of experimental design and timeline for analysis of gene expression in the mPFC of w/t and –e4 mice following context fear recall. *(c)* Freezing during context recall, but not acquisition, is significantly higher in –e4 mice vs. w/t mice (*t*(14) = 2.855, *p* = 0.0127, unpaired t-test). *(d)* Example confocal z-projections from the mPFC showing co-expression of *Slc17a7* (excitatory neurons), *Fos* (recall-activated neurons), and *Ntrk2* (neurons modulated by BDNF signaling). *(e)* A significantly higher proportion of *Ntrk2*/glutamatergic neurons in the PrL co-express recall-induced *Fos* in –e4 mice, while the opposite effect is observed in the IL of –e4 mice (*F*(1,24) = 6.099, *p* = 0.021, interaction for two-way ANOVA). *(f)* Example confocal z-projections from the mPFC showing co-expression of *Gad1* (inhibitory neurons), *Fos,* and *Ntrk2. (g)* In contrast to *Ntrk2*/glutamatergic neurons, *Ntrk2*/GABAergic neurons in the PrL co-express *Fos* at lower levels in –e4 mice following context recall, while *Ntrk2*/GABAergic neurons in the IL co-express recall-induced *Fos* at similar levels in both genotypes (*F*(1,22) = 4.962, *p* = 0.0365, interaction for two-way ANOVA).

Increased recruitment of excitatory neurons in the PrL in -e4 mice during context fear recall suggests that decreased activation of local inhibitory circuits might lead to feed-forward disinhibition of these neurons during exaggerated context fear recall in mutant animals. To explore this possibility, we repeated our experiment with a probe for *Gad1* to assess the degree to which inhibitory neurons in the PrL and IL were recruited during context fear recall in both genotypes (Fig. 4f). We found an opposing phenotype in the PrL of -e4 mice, such that a lower proportion of *Gad1*+/*Ntrk2*+ neurons co-expressed *Fos* in the PrL of -e4 mice compared to w/t mice. These data support the hypothesis that local inhibitory-excitatory micro-circuits in the PrL regulate context fear memory recall, and that these micro-circuits are influenced by BDNF-modulated synaptic input, possibly from the vHC (Fig. 4g). Interestingly, no difference in recruitment of *Gad1*+/*Ntrk2*+ neurons was seen between genotypes in the IL, suggesting that *Ntrk2*-expressing excitatory IL neurons are directly modulated by BDNF released from terminals that originate outside of the IL.

### BDNF-TrkB signaling in vHC-PrL projectors impacts their physiology and recruitment during context fear recall

Our results thus far point to a model in which BDNF synthesized and released from projectors influences plasticity in the PrL to regulate context fear recall. However, it is not known whether vHC-PrL projectors synthesize *Bdnf* during context fear recall, and if so, is fear-related *Bdnf* in projectors derived from promoter IV of the *Bdnf* gene? We again used single-molecule *in situ* hybridization to investigate cell-type specific expression of *tdTomato* (vHC-PrL projectors), *Fos* (recall-activated cells), and exon IV-containing *Bdnf* in ventral CA1 of w/t and –e4 mice following context fear recall (Fig. 5a). We first compared levels of *Fos* expression in projectors between w/t and –e4 mice, and found no significant difference in proportion of projectors co-expressing *Fos* transcripts or number of *Fos* transcripts per projector between genotypes (Supp. Fig. 5a), suggesting that effects of projector activation on downstream brain regions, but not projector activation *per se*, contributes to context fear memory recall. We next compared levels of exon IV-containing *Bdnf* transcripts between vHC-PrL projectors (*tdTomato*+ neurons) and non-projectors (*tdTomato*-neurons) in CA1 in w/t mice, and observed that projectors were significantly enriched for exon IV-containing *Bdnf*, both in proportion of either type of neuron co-expressing exon IV-containing *Bdnf*, and number of exon IV-containing *Bdnf* transcripts per neuron (Supp. Fig. 5b). We further sub-divided our analysis to determine whether projectors that were recruited during context fear recall (*tdTomato*/*Fos*+ neurons) were enriched for exon IV-containing *Bdnf*, and again found that a significantly higher percentage (nearly 100%) of these neurons co-expressed exon IV-containing *Bdnf* (Fig. 5b), and that these neurons expressed a higher number of exon IV-containing *Bdnf* transcripts (Supp. Fig. 5c), as compared to projectors that were not activated during fear recall (*tdTomato*+/*Fos*-) and non-projectors that were activated during fear recall (*tdTomato*-/*Fos*+). While BDNF is not translated from exon IV *Bdnf-Gfp* transcripts in -e4 mice, *Gfp*-containing transcripts can be quantified as a readout of transcriptional activity at promoter IV. We found a similar pattern to w/t in –e4 projectors following context fear memory recall; specifically, projectors activated during recall were highly enriched for exon IV-containing transcripts. Furthermore, a higher percentage of *tdTomato*+/*Fos*+, *tdTomato*+/*Fos*-, and *tdTomato*-/*Fos*+ neurons co-expressed *Bdnf-Gfp* transcripts in -e4 mice following context fear recall, as compared to w/t mice (Fig. 5c). Since it is possible that differences in transcript expression between genotypes, or specificity of the *Bdnf* probe in the -e4 mutants, could drive observed differences in *Bdnf* levels in the vHC, we repeated our experiment with a probe for *Gfp*, and found results that mirrored those observed with *Bdnf* (Supp. Fig. 5d), indicating that transcriptional activity at promoter IV is indeed increased in -e4 mice following context fear recall.

**Figure 5.**
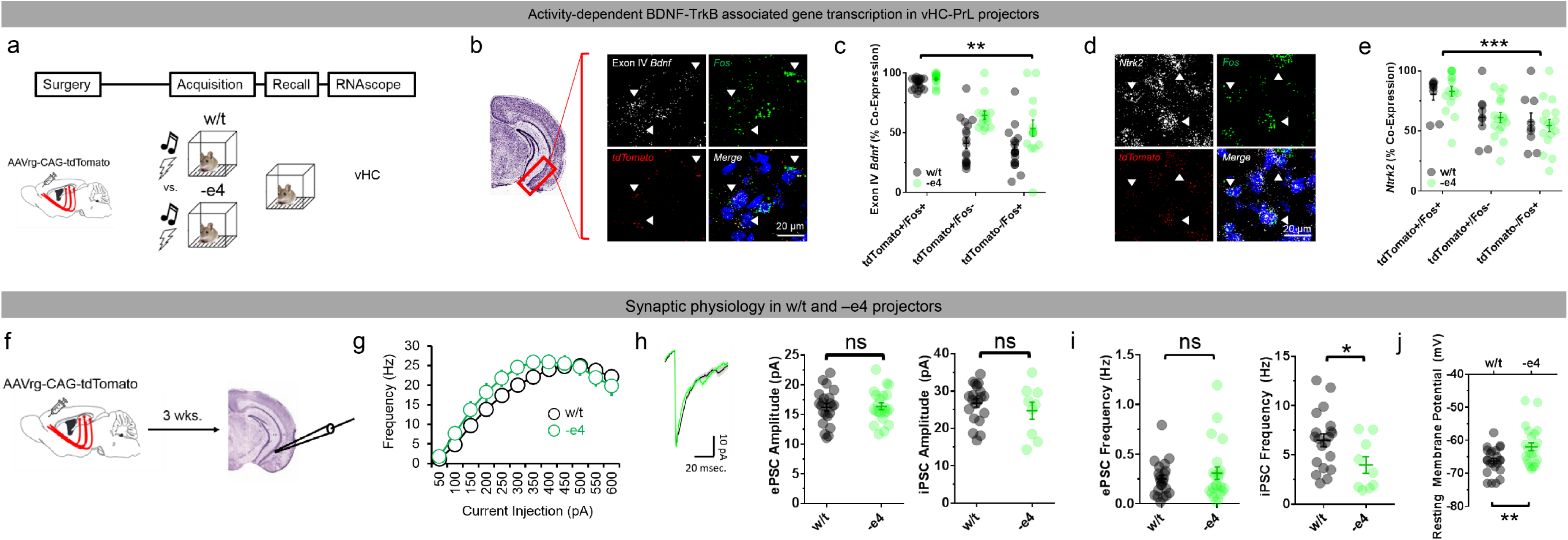
BDNF-TrkB signaling in vHC-PrL projectors impacts their physiology and recruitment during context fear recall. *(a)* Schematic of experimental design and timeline for analysis of gene expression in the vHC of w/t and –e4 mice following context fear recall. *(b)* Example confocal z-projections from ventral CA1 showing co-expression of virally-induced *tdTomato*, context recall-induced *Fos*, and exon IV-containing *Bdnf* mRNA in individual neurons. *(c)* Recall-activated projectors (*tdTomato*+/*Fos*+ neurons) are significantly enriched for exon IV-containing *Bdnf*, as compared to non-activated projectors (*tdTomato*+/*Fos*-) and activated non-projectors (*tdTomato*-/*Fos*+) (*F*(2,60) = 147.6, *p* < 0.0001, main effect of cell type for 2 (genotype) x 3 (cell-type) mixed-factorial ANOVA). *(d)* Example confocal z-projections from ventral CA1 showing co-expression of *tdTomato, Fos,* and *Ntrk2. (e)* Recall-activated projectors are also significantly enriched for *Ntrk2* in both genotypes (*F*(2,46) = 32.57, *p* < 0.0001, main effect of cell type for 2 (genotype) x 3 (cell-type) mixed-factorial ANOVA). *(f)* Schematic of strategy for labeling and recording from vHC-PrL projectors *ex vivo*. *(g)* Input-output curve showing no significant difference in firing rate as a function of current injection between wild/type (w/t) and –e4 mice. *(h)* Neither the amplitude of excitatory (left panel; *t*(42) = 0.17, *p* = 0.87) or inhibitory (right panel; *t*(27) = 0.92, *p* = 0.37, unpaired t-tests) post-synaptic currents (ePSCs and iPSCs, respectively) onto projectors significantly differs between genotypes. *(i)* Frequency of iPSCs (*t*(27) = 2.224, *p* = 0.0347), but not ePSCs (*t*(42) = 0.83, *p* = 0.41, unpaired t-tests), is significantly lower on –e4 projectors. *(j)* Resting membrane potential is significantly more depolarized in –e4 projectors, as compared to w/t projectors (*t*(44) = 2.917, *p* = 0.0055, unpaired t-test).

The observed increase in transcriptional activity at *Bdnf* promoter IV in –e4 vHC-PrL projectors could arise from altered synaptic physiology or excitability of these cells. To test this, we injected AAVrg-CAG-tdTomato into the PrL to label projectors, and cut acute slices of the vHC to record from tdTomato+ cells *ex vivo* (Fig. 5f). We first examined firing rates in vHC-PrL projectors in response to current injection, and found that –e4 projectors had slightly, although non-significantly, higher firing rates after injections of 250 and 300 picoamps (pA), but were otherwise similar to w/t projectors (Fig. 5g), particularly when comparing the maximum number of action potentials in response to current injections (Supp. Fig. 5f). We similarly found that no significant difference in Na^+^ current amplitude (Supp. Fig. 5g) or threshold existed between w/t and –e4 projectors (Supp. Fig. 5h), suggesting that Na^+^ channel-mediated generation of action potentials is intact in these neurons in –e4 animals. We next measured the frequency and amplitude of spontaneous post-synaptic currents (PSCs) onto – e4 and w/t projectors, and found that neither the frequency nor amplitude of excitatory PSCs (ePSCs) onto projectors significantly differed between genotypes (Fig. 5h); however, the frequency of inhibitory PSCs (iPSCs) onto projectors was significantly lower in –e4 mice, even as the amplitude of iPSCs did not significantly differ between w/t and –e4 mice (Fig. 5i). –e4 projectors had a correspondingly more depolarized resting membrane potential (RMP) as compared to w/t projectors (Fig. 5j), in the absence of significant differences in membrane capacitance (Supp. Fig. 5i) or resistance (Supp. Fig. 5j). Taken together, these results suggest that, even though w/t and –e4 projectors generate action potentials at the same rate, action potential generation in –e4 projectors is potentially impacted by altered inhibitory synaptic input.

Given that dendritic release of BDNF from CA1 neurons can trigger autocrine BDNF-TrkB signaling that drives structural and functional plasticity^18^, we also assessed whether vHC-PrL projectors are targets of BDNF signaling by using single-molecule *in situ* hybridization with probes for *Fos, tdTomato,* and *Ntrk2* (Fig. 5d). We found that projectors were significantly enriched for *Ntrk2* mRNA compared to non-projectors in both genotypes (Supp. Fig. 5e), and that projectors activated during context fear recall (*tdTomato*+/*Fos*+) co-expressed *Ntrk2* at the highest levels in both genotypes, with no observed difference in *Ntrk2* co-expression between -e4 and w/t mice (Fig. 5e). Taken together, these results demonstrate that vHC-PrL projectors have distinct plasticity cascades in response to behavior, with projectors being enriched for genes associated with BDNF-TrkB signaling.

### Activation of vHC-PrL projectors produces opposite effects on context fear recall and freezing-related PrL activity in -e4 mice

Given that vHC-PrL projectors are recruited to a similar degree during context fear recall in w/t and -e4 mice, and that –e4 vHC-PrL projectors produce action potentials at the same rate as w/t projectors in response to current stimulation, we reasoned that decreased release of BDNF from vHC-PrL projectors, rather than differences in levels of projector activation *per se*, was driving the observed increases in context fear recall in -e4 mice. In this scenario, activation of BDNF-depleted vHC-PrL projectors in -e4 animals would lead to less pre-synaptic BDNF release and blunted TrkB activation in the PrL, contributing to increases in context fear expression during recall. To test this hypothesis, we virally introduced either the excitatory DREADD receptor hM3Dq or a control reporter (mCherry) into vHC-PrL projectors of -e4 mice, and injected CNO (5 mg/kg, i.p.) 45 m prior to context fear recall (Fig. 6a). In contrast to the suppression of freezing that was observed during context recall following projector activation in w/t mice, projector activation in -e4 mice significantly increased freezing during context recall (Fig. 6b).

**Figure 6.**
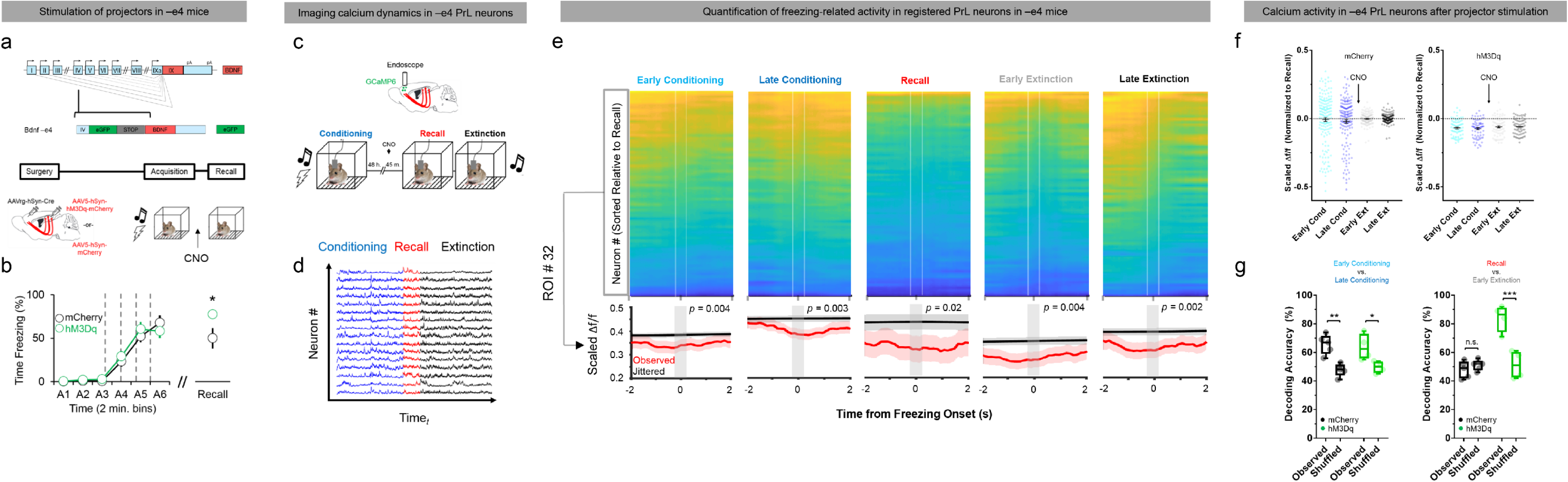
Stimulation of vHC-PrL projectors induces instability in PrL populations during freezing and increases context fear memory expression in –e4 mice. *(a)* Schematic showing viral infection strategy for activation of vHC-PrL projectors prior to context recall in –e4 mice. *(b)* In contrast to w/t mice, activation of projectors significantly increases freezing during context recall in –e4 mice (*t*(15) = 2.365, *p* = 0.03, unpaired t*-*test). *(c)* Cartoon of strategy for imaging calcium activity in PrL neurons following projector activation in –e4 mice. *(d)* Example calcium traces from 15 simultaneously imaged PrL neurons in an –e4 mouse registered across conditioning, context recall, and extinction training. *(e) (Top row)* Averaged activity from 66 simultaneously imaged PrL neurons from one –e4 mCherry mouse across early conditioning (pre-shock), late conditioning (post-shock), context recall, early extinction, and late extinction during freezing bouts. Neurons are sorted relative to their Δf/f value during context recall freezing onset (dashed white lines). PrL population activity patterns during freezing remain similar between fear memory phases. *(Bottom row)* Observed and jittered (peri-freezing) activity traces from one neuron in the population. This neuron shows the same pattern of freezing-related activity during all fear memory phases. *(f)* On average, calcium activity during freezing bouts does not significantly differ between any fear memory phase and context recall (black dashed line at 0 on y-axis) in mCherry control –e4 mice (left panel). In hM3Dq mice, CNO injections caused calcium activity during context recall to become significantly higher relative to calcium activity during other fear memory phases (black dashed line, right panel; *F*(4) = 4.52, *p* = 0.001, significant group x fear memory phase interaction for linear mixed effects model). *(g)* Decoding accuracy of a linear classifier using freezing-related calcium activity to predict fear memory phase (observed) is significantly above chance (shuffled data) when distinguishing between early and late conditioning for both groups (left panel; mCherry *p* = 0.002, hM3Dq *p* = 0.012), but falls to chance levels when distinguishing between context recall and early extinction in the mCherry group (right panel; mCherry *p* = 0.74, hM3Dq *p* = 0.0004, Sidak’s multiple comparison tests).

We next hypothesized that this increase in fear recall could be driven by differences in how projector stimulation impacted population dynamics in PrL neurons of -e4 mice. To test this hypothesis, we compared calcium activity in the PrL of mCherry controls in which projectors were left un-stimulated to hM3Dq mice in which projectors were stimulated prior to context recall (Fig. 6c & 6d). In mCherry -e4 control mice, individual PrL neurons tended to maintain a consistent freezing-related activity profile across fear memory phases (Fig. 6e), suggesting that abnormal context recall in -e4 mice may be due to stability in population representations between fear epochs. To test this possibility, we used a linear mixed effects model to quantify differences in how PrL neurons represent freezing during fear memory phases, and found that freezing-related calcium activity did not significantly differ from context recall for any fear memory phase in the mCherry group. However, our model did reveal a significant interaction between fear memory phase and stimulation group in predicting freezing-related calcium activity, with the interaction driven by differences in freezing-related calcium activity between context recall and other fear memory phases in hM3Dq animals (Fig. 6f). These results suggest that projector stimulation in –e4 mice paradoxically destabilizes freezing-related calcium activity during context recall.

To determine whether freezing-related PrL calcium activity can predict fear memory phase in –e4 animals, we again used a SVM linear classifier to predict whether a freezing bout occurred during early and late conditioning, and context recall and extinction training. The classifier discriminated between early and late conditioning freezing bouts at levels above chance for both groups, but failed to distinguish between context recall and extinction training only in the mCherry group (Fig. 6g). These results demonstrate that decreased activity-dependent BDNF production impedes the ability of PrL neurons to dynamically represent distinct fear memory phases, and that stimulation of vHC-PrL projectors in the absence of promoter IV-derived BDNF paradoxically increases freezing during context fear recall, potentially by decreasing PrL ensemble stability.

### Expression of BDNF selectively in vHC-PrL projectors reverses behavioral and molecular phenotypes during context fear recall in -e4 mice

Our data thus far support a model in which activation of vHC-PrL projectors induces release of promoter IV-derived BDNF into the PrL to modify excitatory and inhibitory post-synaptic targets to drive suppression of contextually-mediated fear memory recall. If decreased BDNF production and release from vHC-PrL projectors increases context fear recall, we reasoned that exogenous expression of BDNF in vHC-PrL projectors would be sufficient to decrease exaggerated context fear recall in - e4 animals. To test this hypothesis, we used a combination of viruses to selectively express BDNF in vHC-PrL projectors of -e4 mice. Specifically, we injected AAVrg-hSyn1-emBFP-Cre into the PrL and AAV8-CAG-FLEX-BDNF:HA into the vHC for the Cre-dependent expression of a BDNF expression cassette in our experimental group, and injected AAV8-hSyn1-DIO-mCherry into the vHC in our control group for Cre-dependent expression of the mCherry reporter (Fig. 7a). In our construct, BDNF is fused to a hemagglutinin (HA) tag, which was readily detectable in vHC-PrL projectors with anti-HA immunohistochemistry (Fig. 7b). Six weeks following virus injections, both groups were fear conditioned and tested for context recall. We found that selective BDNF expression in projectors (BDNF:HA group) decreased freezing during recall (Fig. 7c), supporting our hypothesis that BDNF synthesis in projectors is sufficient to suppress context fear.

**Figure 7.**
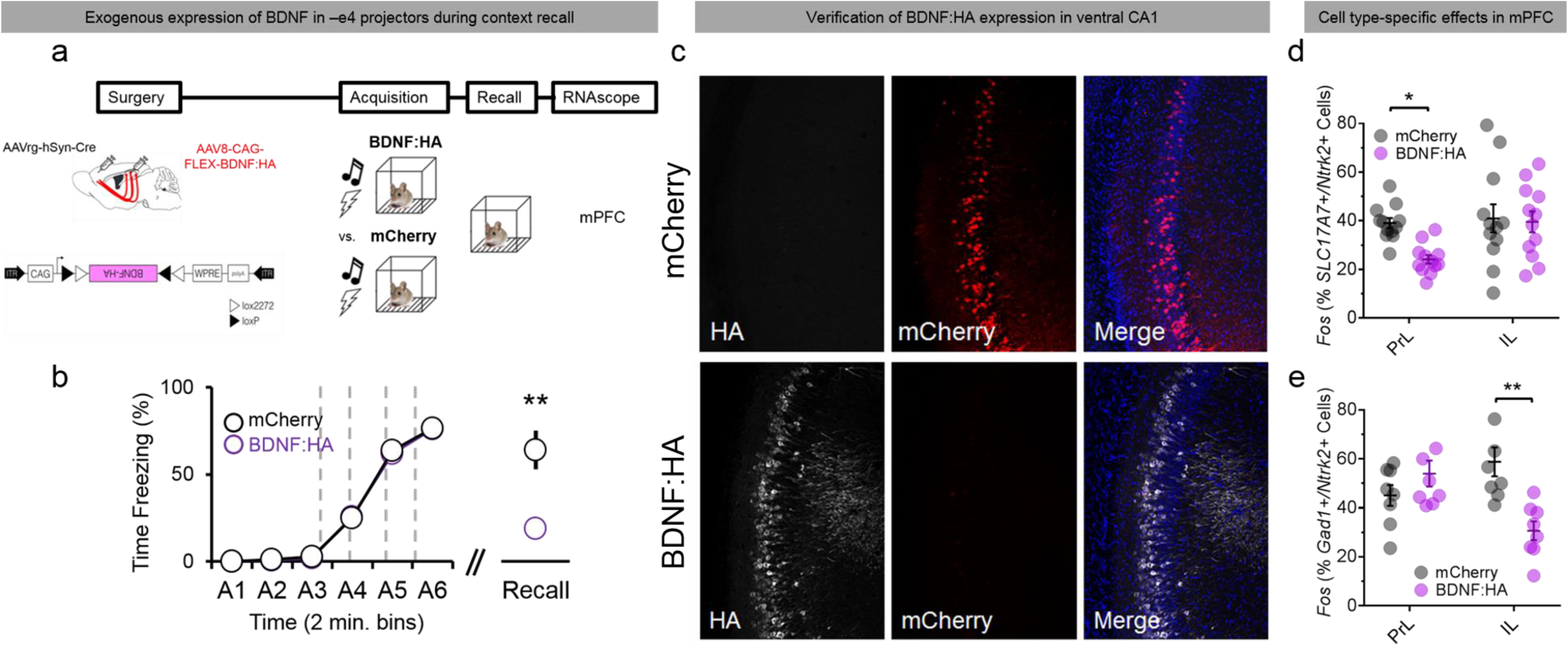
Exogenous expression of BDNF protein in vHC-PrL projectors reverses fear-related behavioral and molecular phenotypes in –e4 mice. *(a)* Schematic of timeline and experimental design for viral over-expression of BDNF in –e4 projectors. Bottom panel shows sequence for AAV8-CAG-FLEX-BDNF:HA construct. *(b)* Over-expression of BDNF in projectors reduces freezing in –e4 mice during context recall, but not fear acquisition (*t*(14) = 3.753, *p* = 0.0021, unpaired t-test). *(c)* Example confocal z-projections from ventral CA1 showing mCherry expression in control –e4 animals, and immunofluorescence for HA protein in BDNF:HA animals. *(d)* BDNF over-expression in –e4 projectors reduces the proportion of *Ntrk2*/glutamatergic neurons that co-express *Fos* in the PrL (*F*(1,22) = 5.423, *p* = 0.0295, interaction for two-way ANOVA). *(e)* BDNF over-expression in –e4 projectors reduces the proportion of *Ntrk2*/GABAergic neurons that co-express *Fos* in the IL (*F*(1,14) = 13.7, *p* = 0.0024, interaction for two-way ANOVA).

Ninety minutes after context fear recall, we sacrificed mice in both groups and used single-molecule *in situ* hybridization to measure effects of exogenous BDNF expression on recruitment of BDNF-receiving neurons in the PrL and IL. We found that the proportion of *Slc17a7*+/*Ntrk2*+ neurons that co-expressed fear-induced *Fos* was significantly lower in the PrL of BDNF:HA mice, providing evidence that BDNF release from projectors modifies post-synaptic signaling in excitatory neurons in the PrL to suppress context fear recall. Importantly, we did not find any significant difference in recruitment of *Slc17a7*+/*Ntk2*+ neurons between groups in the IL, indicating that changes in IL function during context fear recall in -e4 mice are not driven by differences in PrL signaling (Fig. 7d). Conversely, although the proportion of *Gad1*+/*Ntrk2*+ neurons co-expressing *Fos* was higher in the PrL of BDNF:HA mice, this effect was not statistically significant. One possible explanation for this result is that a sub-population of GABergic interneurons may regulate inhibitory signaling onto excitatory neurons in the PrL following BDNF release from projectors, an effect that could be obscured when using *Gad1* to broadly identify all inhibitory neurons in this brain area. Unexpectedly, we did find that *Fos* co-expression in *Gad1*+/*Ntrk2*+ neurons was significantly lower in the IL of BDNF:HA mice (Fig. 7e). This result suggests that decreased recruitment of PrL excitatory neurons during context fear suppression may directly influence inhibitory signaling in the IL, possibly via direct synaptic contact between excitatory PrL neurons and inhibitory IL neurons. Given that we did not observe an opposite phenotype in -e4 mice during context fear recall (i.e., *increased* recruitment of inhibitory neurons in the IL), this suggests that parallel pathways to the PrL and IL likely contribute to increased fear expression in mutant animals, and that restoring function in one pathway (vHC-PrL projections) is sufficient to overcome dysfunction in other pathways during contextually-mediated fear recall.

## Discussion

We provide evidence that vHC-PrL projection neurons are a molecularly distinct sub-population in the vHC that is enriched for gene sets implicated in learning, memory, and stress signaling pathways and associated with neuropsychiatric disorders that feature fear dysregulation (PTSD and GAD). Activation of vHC-PrL projection neurons suppresses context fear memory recall and inhibits flexible coding of distinct contextually-influenced fear states in PrL neurons. Activation of these projection neurons also induces unique patterns of activity-dependent gene expression in the mPFC, which largely resemble gene expression patterns in cultured neurons after BDNF application. Mice with impaired activity-dependent BDNF expression (-e4 mice) demonstrate increased freezing during context fear recall, as well as aberrant inhibitory signaling onto vHC-PrL projectors. These mutant mice also have impaired recruitment of excitatory and inhibitory *Ntrk2*-expressing neurons in the PrL following context fear recall. Activation of vHC-PrL projectors in –e4 mice causes opposite behavioral (increased freezing) and neuronal (unstable representations of fear memory phase) phenotypes compared to w/t mice, suggesting that activation of vHC inputs can have divergent effects on PrL-associated behavior and neuronal activity as a function of which molecules are synthesized and released from those inputs. Finally, we demonstrate that exogenous expression of BDNF in –e4 projectors reverses behavioral and molecular phenotypes in the PrL during context fear recall.

### vHC-PrL Projectors Alter PrL Population Dynamics to Suppress Context Fear Memory Recall

Our findings demonstrate a novel role for direct ventral hippocampal inputs to the PrL in suppression of freezing during recall of a fear-associated context. The vHC is known to be necessary for both tone^19, 20^ and context^14^ fear retrieval, but the efferent projections within the vHC that mediate these processes are not known. Early studies showed that vHC inactivation increases firing rates in PrL neurons during tone presentations in conditioned animals^21^, suggesting that vHC outputs to the PrL might mediate retrieval of tone-shock associations. The use of muscimol to inactivate vHC neurons in that study, however, precluded a conclusion about how PrL-projecting neurons in the vHC participate in this process. Indeed, a recent study in which DREADD receptors were expressed exclusively in vHC-PrL projectors revealed that neither activation nor inactivation of these neurons affects freezing during tone retrieval in conditioned mice^22^. This same study reported that activation of this pathway decreases freezing during contextually-dependent renewal of fear following extinction, suggesting that vHC-PrL projectors may generally support the retrieval of context-fear associations. Inactivation of the PrL decreases freezing during both tone^7^ and context^11^ recall, while micro-stimulation of the PrL increases freezing during tone presentations in conditioned animals^23^. In contrast, local inactivation of the IL does not affect freezing during context recall, but does impair extinction learning and extinction recall^8^. These results support the canonical view that the PrL and IL play doubly dissociable roles in fear retrieval and extinction; specifically, that the PrL is critical for retrieval (increased freezing), but not extinction (decreased freezing), and vice versa for the IL.

Recent studies, however, have called into question the view that the PrL supports increased fear expression (freezing). Activation of a subset of PrL neurons that project to the IL causes decreased freezing (facilitates extinction)^9^, demonstrating that heterogeneous populations of neurons in the PrL can control distinct aspects of fear behavior. Indeed, largely non-overlapping sub-populations of neurons in the PrL are active during fear memory formation and retrieval; furthermore, forced depolarization of PrL neurons that were previously active during fear memory acquisition is not sufficient to elicit freezing 24 h later^24^, indicating that disparate PrL ensembles dynamically represent temporally-distinct fear memory phases. Our data extend these results by showing that PrL ensembles that are active across fear memory phases also dynamically distinguish between them, representing freezing behavior differently as a function of fear memory acquisition, retrieval, or extinction. Moreover, freezing bouts during distinct fear memory phases are best represented by shifts in activity patterns within PrL populations, as evidenced by the ability of population activity to predict in which fear memory phase the animal is freezing (Fig. 2h). These data complement other studies showing that PFC neurons exhibit “mixed selectivity” for a variety of task parameters depending on the behavioral context; e.g., PFC neurons in non-human primates that respond to both spatial location and object identity^25^, and mPFC neurons in mice that respond to sensory, motor, and response outcome variables during goal-directed behavior^26^. We show that activation of vHC inputs to the PrL renders PrL populations less flexible between context fear memory retrieval and extinction, indicating that the vHC affects freezing behavior during retrieval in part by affecting the ability of PrL neurons to distinguish between behaviorally-relevant contexts.

### Molecular Signatures in the vHC-PrL Pathway

We demonstrate that, in addition to sharing common efferent projection targets, vHC-PrL projectors are molecularly distinct from other vHC neurons (Fig. 1c), including those in CA1 that do not project to the PrL (Fig. 5c & Fig. 5e). These results indicate that genetically distinct sub-populations in the vHC^27^ may send axons that synapse onto similar downstream brain areas. We further show that vHC-PrL projectors are enriched for gene sets that are implicated in GAD and PTSD, suggesting that molecular profiles of projection-specific populations in rodents could be “mapped on” to risk gene sets in human neuropsychiatric disorders. Synthetic activation of vHC-PrL projectors induced unique activity-dependent gene programs in the mPFC, further demonstrating that the mPFC transcriptome can be dynamically affected by activation of anatomically-distinct upstream inputs. Activity-dependent gene expression profiles differ according to an animal’s experience in a number of different brain areas, including the mPFC^28^, indicating that distinct events (i.e., fear memory retrieval vs. social interaction) can have unique experience-dependent “molecular signatures”^29^. The identification of transcriptomic signatures related to activation of spatially-localized neural circuits could compliment these findings by linking these circuits with defined behaviors at the molecular level. These findings may also have implications for the identification of unique biomarkers associated with neuropsychiatric disorders, many of which are associated with aberrant PFC function^30^.

### Activity-Dependent BDNF Signaling Alters vHC-PrL Function During Context Fear Memory Recall

Our RNA-seq data revealed that genes associated with BDNF-TrkB signaling are broadly up-regulated in the mPFC in response to stimulation of vHC inputs. Specifically, we demonstrate that fold-changes in our RNA-seq data set are highly correlated with fold-changes in another RNA-seq data set of cultured neurons following application of BDNF^17^, indicating that stimulation of vHC-PrL projectors induces BDNF-TrkB signaling within the mPFC. A host of previous work has demonstrated relationships between hippocampal-prefrontal function, BDNF, and fear behavior^15, 16^, including work from our laboratory showing that mice with decreased production of activity-dependent BDNF have deficits in fear extinction that co-occur with impaired hippocampal-prefrontal oscillatory synchrony^31^. Transcription from promoter IV of the *Bdnf* gene in the dorsal hippocampus is correlated with fear memory consolidation^32^ and epigenetic modification of promoter IV in the IL is linked with fear extinction^33, 34^, findings that suggest that BDNF production from promoter IV may contribute to the mechanism underlying fear behavior^35^. Our lab recently demonstrated that synthetic excitation of cells in the ventral dentate gyrus expressing promoter IV-derived *Bdnf* transcripts increased freezing during context fear retrieval and concomitantly attenuated vHC-PrL oscillatory synchrony^36^. The current data extend this finding by showing that promoter IV-derived *Bdnf* is enriched in a projection-specific population of vHC neurons and that global disruption of promoter IV-derived BDNF directly impacts function in this projector population.

Our results suggest a dual mechanism supporting context fear suppression in w/t animals in which release of BDNF from vHC-PrL projectors in response to their excitation silences a subset of excitatory neurons in the PrL, and excites a subset of excitatory neurons in the IL. *Ntrk2*+ excitatory neurons in the IL may be modulated during context fear expression by BDNF from a number of brain areas synapsing directly onto IL neurons, including the vHC. It is also possible that these IL neurons are sequentially activated by *Bdnf*-synthesizing PrL neurons following stimulation of inputs from the vHC. This demonstrates that pre-synaptic release of BDNF from vHC-PrL projectors, which are enriched for exon IV-containing *Bdnf*, likely modulates context fear recall by regulating excitatory/inhibitory function in the post-synaptic PrL.

Our data show that PrL population activity during freezing is more rigid between fear memory phases in –e4 animals, as compared to w/t animals. This suggests that increased fear expression during context recall in these mice may be driven by abnormal stability in PrL coding between distinct fear epochs. Depolarization of vHC-PrL projectors in –e4 mice additionally led to dissociable effects on behavior and population activity in the PrL during context fear recall. Paradoxically, in contrast to w/t animals, stimulation of vHC-PrL projectors increased freezing during context fear recall, suggesting that synthesis and release of activity-dependent BDNF from projectors, and not activation of projectors *per se*, drives decreased contextually-driven fear expression. vHC-PrL projector stimulation also paradoxically caused PrL neurons to differentially represent freezing bouts during context recall and extinction training, indicating that molecule by circuit interactions in the vHC-PrL pathway critically govern behavior during recall of context-fear memories.

We found that exogenous expression of BDNF protein selectively in vHC-PrL projectors reversed behavioral and molecular phenotypes in the PrL of –e4 mice during context fear recall. This finding demonstrates that manipulation of molecular function within spatially-defined circuits can impact fear-related behavior, even in animals with developmental alterations in brain function. These data demonstrate that genetic background can critically bias circuit function, which must be taken into account for circuit-targeting strategies in symptoms of neuropsychiatric disease; for example, in individuals with functional single-nucleotide polymorphisms (SNPs) in the *BDNF* gene, which can affect activity-dependent BDNF secretion^40^ and fear behavior^41^.

### Summary

Our findings support a model in which activity-dependent release of BDNF from vHC neurons modulates post-synaptic signaling in both inhibitory and excitatory PrL neurons, resulting in modification of contextual representations in PrL populations to suppress freezing during fear memory retrieval. We also show that vHC-PrL projectors are a molecularly distinct sub-class of vHC neurons, and that a number of genes that are differentially-expressed in projectors are also implicated in PTSD and GAD. Our data add to a growing number of studies elucidating the circuitry underlying fear behavior with projection specificity – for example, IL and BLA-projecting vHC neurons in context-dependent renewal of fear following extinction^42, 43^, and ventral midline thalamic-projecting mPFC neurons in fear extinction^44^. We further causally link molecular function in this projection-specific population with context fear recall, expanding on research showing that manipulating BDNF in either the vHC or mPFC affects extinction of conditioned fear^15^ and avoidance behavior^45^ by directly linking BDNF-TrkB signaling within a projection-specific sub-population with contextually-dependent fear behavior. Taken together, our data provide mechanistic insight into how hippocampal-prefrontal circuitry regulates context fear memory expression at the molecular, cellular, and systems levels, information which will be critical for informing eventual precision medicine approaches that target fear-related symptoms in disorders such as PTSD and GAD.

## Methods

### Subjects

Mice with decreased production of activity-dependent BDNF (-e4) mice were generated as previously described^20^. Briefly, an enhanced green fluorescent protein-STOP cassette was inserted upstream of promoter IV of the *Bdnf* gene, so that transcription initiated from promoter IV produces green fluorescent protein *in lieu* of BDNF. Male wild-type (w/t) C57Bl6/J and -e4 mice were group-housed (3-5 animals per cage) and maintained on a 12 h light/dark cycle in a temperature and humidity-controlled colony room. All animals had *ad libitum* access to food and water. Animals were 10-16 weeks of age, and weighed 25-35 g at time of surgery. All procedures were in accordance with the Institutional Animal Care and Use Committee of SoBran Biosciences Inc.

### Surgical Procedures

Mice were anesthetized with isoflurane (1-2.5% in oxygen), placed into a stereotaxic frame, and an incision was made along the midline of the scalp. The skull was leveled, bregma was identified, and small holes were drilled with a 0.9mm burr (Fine Science Tools) above the vHC (−3.2mm AP, ±3.1mm ML, −3.4mm DV) and PrL (+1.7mm AP, ±0.3mm ML, −1.7mm DV) for viral injections. For c-Fos immunohistochemistry, *in situ* hybridization, and *ex vivo* electrophysiology experiments, AAVrg-CAG-tdTomato (7.2×10^12 gc/ml) was used for injection into the mPFC. For calcium imaging experiments, 5 µl of AAV-Syn-GCaMP6f-WPRE-8V40 (20 ng/µl) was mixed with 5 µl of AAVrg-hSyn-Cre-WPRE-hGH (1.2×10^13 gc/ml) for injection into the PrL and AAV5-hSyn-DIO-hM3Dq-mCherry (3.8×10^12 virus molecules/ml) or AAV5-hSyn-DIO-mCherry (4.8×10^12 gc/ml) were used for injection into the vHC. For experiments wherein vHC-PrL projectors were manipulated by DREADD receptors, AAVrg-hSyn-Cre-WPRE-hGH (1.2×10^13 gc/ml) was used for injection into the PrL and AAV5-hSyn-DIO-hM3Dq-mCherry (3.8×10^12 virus molecules/ml) or AAV5-hSyn-DIO-mCherry (4.8×10^12 gc/ml) were used for injection into the vHC. For experiments involving exogenous expression of BDNF in vHC-PrL projectors, AAVrg-hSyn-Cre-WPRE-hGH (1.2×10^13 gc/ml) was used for injection into the mPFC and AAV8-CAG-FLEX-BDNF:HA (1.4×10^13 gc/ml) was used for injection into the vHC. For all infusions, a total volume of 600 nl/hemisphere was injected into the vHC or PrL at a rate of 200 nl/min via a 10 µl syringe (Hamilton) controlled by an automated infusion pump (World Precision Instruments). Mice received a sub-cutaneous injection of a local anesthetic (Bupivicaine) along the incision site, as well as an i.p. injection of an analgesic (Meloxicam, 5 mg/kg) during surgery. Meloxicam injections were given for three days post-surgery for pain relief.

### Behavior

For fear conditioning, animals were conditioned to associate a 30 s tone (4000 Hz) with a 2 s, 0.6 mA foot-shock. 4 tone-shock combinations were given, with the shock co-terminating with the last 2 s of each tone presentation. Tone-shock combinations were separated by 90 s intervals. 48 h later, mice were placed back into the conditioning chamber to measure recall of the conditioning context (in the absence of the tone) for 180 s. For calcium imaging experiments, a total of 23 tones (30 s each) were presented in the absence of the foot-shock for tone/context extinction training following pre-tone context recall. Tones were separated by 5 s intervals. Freezing (cessation of movement) during acquisition and recall sessions was quantified with automated tracking software (FreezeScan; CleverSys, Inc.). For calcium imaging experiments, the animal’s body position was extracted per frame (30 fps) with an automated mouse tracking package in MATLAB (motr; Janelia Farms^46^). Movement between frames was calculated by taking the Euclidean distance between body coordinates from frame to frame. Freezing bouts for calcium imaging experiments were defined to start at the frame in which a mouse showed no movement following a frame in which it had shown movement. For mice with either hM3Dq or mCherry expression, clozapine-N-oxide (CNO, Tocris; 5 mg/kg, i.p.) dissolved in 1x PBS was administered 45 min prior to context recall. For mice used to measure c-Fos expression in the vHC following fear recall using immunohistochemistry, half of the mice were given foot-shocks during acquisition, and half of the mice were presented with the tone in the absence of a foot-shock.

### Immunohistochemistry (c-Fos and HA)

For c-Fos immunohistochemistry, mice were trans-cardially perfused with 4% paraformaldehyde 2 h following the termination of context recall. Brains were extracted and stored in 4% paraformaldehyde for 24 h, and cryoprotected in 30% sucrose in 1x PBS/sodium azide (0.05%) for 2-3 d. Coronal sections (50 μm) of the vHC were cut on a sliding microtome (Leica) with attached freezing stage (Physitemp), washed in 5% Tween-80 in 1x PBS, and incubated in blocking solution (0.5% Tween-80, 5% normal goat serum in 1x PBS) with agitation for 6-8 h. The sections were then incubated in 1:1000 anti-Fos antibody (Millipore; cat # ABE457) in blocking solution overnight at 4°C with agitation. The following day, the sections were washed, incubated in 1:1000 goat anti-rabbit AlexaFluor 647 (Sigma) in blocking solution for 2 h with agitation, washed again, and incubated in 1:5000 DAPI (Sigma) in 1x PBS for 20 min. The sections were then mounted and coverslipped, and co-expression of tdTomato/c-Fos (for analysis of recruitment of vHC-PrL projectors cells during context fear recall) was visualized at 20x magnification on a Zeiss 700 LSM confocal microscope.

For HA immunohistochemistry, vHC slices from either Rpl22^HA^/RiboTag mice or mice with expression of AAV8.CAG.FLEX.BDNF:HA were washed in 1x PBS, permeabilized for 20 min in 0.5% Triton X-100 in 1xPBS, and incubated in blocking solution (0.25% Triton X-100, 5% normal goat serum in 1x PBS) with agitation for 1 h. The sections were then incubated in 1:400 anti-HA antibody (Sigma; cat # H6908) in blocking solution for 48 h at 4°C with agitation. Next, the sections were washed, incubated in 1:500 goat anti-rabbit AlexaFluor 647 (Sigma) in blocking solution for 2 h with agitation, washed again, and incubated in 1:5000 DAPI (Sigma) in 1x PBS for 20 min. The sections were then mounted and coverslipped, and expression of HA was visualized at 20x magnification on a Zeiss 700 LSM confocal microscope. c-Fos, tdTomato, mCherry, and HA-expressing cells were quantified from maximum intensity projections created from z-stacked, tiled images.

### Single-Molecule *In Situ* Hybridization

For analysis of cell type-specific gene expression following context fear recall, animals were killed 90 m following context recall and brains were immediately extracted and flash-frozen in 2-methylbutane (ThermoFisher). Coronal sections of vHC and mPFC (16 µm) were cut, and mounted onto slides (VWR, SuperFrost Plus). The slides were quickly fixed in 10% buffered formalin at 4°C, washed in 1X PBS, and dehydrated in ethanol. Slides were next pre-treated with a protease solution, and subsequently incubated at 40°C for 2 h in a HybEZ oven (ACDBio) with a combination of probes for relevant mRNA^47^. Following incubation with the probes, the slides were incubated at 40°C with a series of fluorescent amplification buffers. DAPI was applied to the slides, and the slides were cover-slipped with Fluoro-Gold (SouthernBiotech). Transcript expression was visualized on a Zeiss LSM 700 confocal microscope with a 40x oil-immersion lens (exon IV-containing *Bdnf*- 647 nm, *tdTomato*- 555 nm, *Fos*- 488 nm, DAPI- 405 nm for analysis of exon IV-containing *Bdnf* in vHC-PrL projectors; *Fos*- 647 nm, *tdTomato*- 555 nm, *Ntrk2*- 488 nm, DAPI- 405 nm for analysis of *Ntrk2* in vHC-PrL projectors; *SLC17A7*- 647 nm, *Fos*- 555 nm, *Ntrk2*- 488 nm, DAPI- 405 nm for analysis of recruitment of excitatory *Ntrk2*-expressing neurons in the mPFC; *Gad1*- 647 nm, *Fos*- 555 nm, *Ntrk2*- 488 nm, DAPI- 405 nm for analysis of recruitment of inhibitory *Ntrk2*-expressing neurons in the mPFC). For quantification of transcript co-localization, 40x z-stacks from ventral CA1, the PrL, and IL were taken, and custom MATLAB functions were used to first isolate cell nuclei from the DAPI channel using the cellsegm toolbox^48^ combined with a watershed algorithm for cell splitting in 3 dimensions. Once centers and boundaries of individual cells were isolated, an intensity threshold was set for transcript detection, and watershed analysis was used to split the remaining pixels in each channel into identified transcripts. Custom MATLAB functions^49^ were then used to determine the size of each detected transcript (*regionprops3* function in Image Processing toolbox), and split unusually large areas of fluorescence into multiple transcripts based on known transcript size. Each transcript was then assigned to a cell based on its position in 3 dimensions; transcripts that were detected outside of the boundaries of a cell were excluded from further analysis.

### Calcium Imaging

For calcium imaging experiments, an endoscopic lens (1mm diameter, 4mm length; ProView, Inscopix) targeting the PrL (+1.7mm AP, ±0.3mm ML, −1.4mm DV) was implanted 2 weeks following virus injections. The lens was attached to 3 small skull screws (Fine Science Tools) with dental acrylic (Lang Dental). Black dental acrylic (Lang Dental) was then built up around the lens to obscure outside light. The top of the lens was protected with KwikCast (World Precision Instruments). 4-6 weeks following lens implantation, a baseplate (Inscopix) was attached with black dental acrylic to interface with a one-photon tethered miniscope (nVista 2.0; Inscopix) during fear conditioning. Neuronal activity in the PrL was imaged and recorded with nVista HD software (Inscopix; 30 fps) during fear memory acquisition, context recall, and extinction training. Movies were temporally down-sampled to 15 fps with Inscopix Data Processing Software, and motion-corrected with NoRMCorre in MATLAB^50^. Next, position of and temporal fluorescent traces from ROIs (cells) were extracted and longitudinally registered across fear phases within animals with the CNMF-E toolbox in MATLAB^51^. Traces were visually inspected for noise artifacts, and normalized to values between 0 and 1. For classifying cells into the three activity groups of “freezing-excited”, “freezing-inhibited”, and “freezing-neutral”, a distribution of the cell’s observed calcium activity during freezing bouts (from 1 s prior and 1 s following freezing onset) was compared with a distribution of calcium activity from that same cell permuted around the freezing bout (from −2 s to −1 s prior, and +1 to +2 s following freezing onset) in 200 ms shifts. Wilcoxon’s rank-sum tests were used to compare calcium activity during freezing bouts (observed) to permuted activity (jittered) for each cell. If observed calcium activity was greater than jittered activity with a significant *p* value, that neuron was categorized as “freezing-excited”. If the inverse was true, the neuron was categorized as “freezing-inhibited”. If the *p* value for the rank-sum test was > 0.05, that cell was categorized as “freezing-neutral”.

### Analysis of Population Activity in the PrL During Freezing

To compare freezing-related calcium activity across fear memory phases between groups (mCherry vs. hM3Dq), we used a linear mixed effects model with scaled Δf/f as the dependent variable and main effects for stimulation group, fear memory phase, and their statistical interaction, with random intercepts of mouse and neuron. We fit these models using the R lmerTest package^52^, corresponding to the following code: ‘lmer(y ∼ Condition*Manipulation + (1|Mouse) + (1|Neuron)’. For PCA, average scaled calcium activity per cell across freezing bouts (−1 and +1 s surrounding freezing onset) was calculated for all animals in the mCherry group for each fear phase (early conditioning, late conditioning, recall, early extinction, and late extinction), and used as input into a 225 (45 bins*5 fear phases = 225) x 230 (number of registered cells across all mCherry animals). For machine learning-based classification of fear phase, we used a support vector machine (SVM) linear classifier to distinguish between a) early conditioning and late conditioning, and b) recall and early extinction training based on PrL population activity during freezing bouts (−1 and +1 s surrounding freezing onset). We categorized each freezing bout according to the fear phase in which it occurred, and used these categories as training labels for the classifier (e.g., 0 for early conditioning, 1 for late conditioning). We then used a “leave one out” cross-validation approach^53, 54^ in which we selected one freezing bout for use as a testing set, trained the classifier on all remaining freezing bouts, and repeated this procedure until all freezing bouts had been used as a testing set. We then calculated decoding accuracy by comparing the classifier output with the training label (either 0% correct or 100% correct), and then averaged decoding accuracy from all freezing bouts for each animal. For all animals, the classifier’s penalty parameter was set to “1”. Decoding accuracy was calculated with custom MATLAB functions, and with the “libsvm”^55^ toolbox in MATLAB.

### Ex Vivo Electrophysiology

To patch onto vHC-PrL projectors in w/t and –e4 animals, mice were killed following isoflurane administration, and brains were quickly removed. 300 μm thick slices of the vHC were cut on a Leica VT1000 S Vibrating blade microtome (Leica Biosystems, Inc.). Slices were maintained in oxygenated ice-cold Na^+^-free sucrose solution, and initially incubated at 34°C in a Ringer solution (ACSF) containing 125 mM NaCl, 2.5 mM KCl, 125 mM NaH_2_PO_4_, 2 mM CaCl_2_, 1 mM MgCl_2_, 26 mM NaHCO_3_, and 10 mM dextrose. Slices were equilibrated for at least 30 m before recording. For current clamp, recording pipettes were filled with intracellular solution containing 130 mM K-gluconate, 1 mM MgCl_2_, 5 mM EGTA, 5 mM MgATP, 10 mM HEPES, and 0.4 mM Na_2_GTP. tdTomato+ cells were imaged on an Olympus Bx51W1 microscope. Resistances of patch pipettes were 4-7 MΩ. Signals were amplified and filtered at 2 kHz with Axopatch 200B (Molecular Devices, Sunnyvale, CA) and acquired at sampling intervals of 20-100 μs through a DigiData 1321A interface with pCLAMP 10 (Molecular Devices, Sunnyvale, CA). The access resistance was monitored during recordings, and the data were excluded from analysis if the series resistance changed more than 20% from control levels (10-25 MΩ). Frequency and amplitude of individual ePSCs and iPSCs were analyzed with Clampfit 10 (Molecular Devices). Input resistances were calculated offline from the voltage produced by negative current injection (−20 pA) prior to the step currents.

### qPCR and Bulk RNA Sequencing

For analysis of activity-dependent gene expression in the mPFC following DREADD (hM3Dq) activation in projectors, we gave either CNO (5 mg/kg, i.p.) or vehicle (saline) injections and killed animals 135 m later. Brains were extracted, and the mPFC was dissected out and flash frozen in 2-methylbutane (Fisher). Total RNA was isolated and extracted from the tissue using TRIzol (Life Technologies, Carlsbad, CA), purified using RNeasy minicolumns (Qiagen, Valencia, CA), and quantified using a NanoDrop spectrophotometer (Agilent Technologies, Savage, MD). For qPCR, RNA concentration was normalized and reverse transcribed into single-stranded cDNA using Superscript III (Life Technologies). qPCR was performed using a Realplex thermocycler (Eppendorf, Hamburg, Germany) with GEMM mastermix (Life Technologies) with 40 ng of synthesized cDNA. Individual mRNA levels for *Fos, Arc,* and *Npas4* were normalized for each well to *Gapdh* mRNA levels. For RNA-seq, the Nextera XT DNA Library Preparation Kit was used to generate sequencing libraries according to manufacturer instructions, and samples were sequenced on a HiSeq2000 (Illumina). Reads were aligned to the mm10 genome using the HISAT2 splice-aware aligner^56^, and alignments overlapping genes were counted using featureCounts version 1.5.0-p3 relative to Gencode version M11. Differential expression analyses were performed on gene counts using the voom approach^57^ in the limma R/Bioconductor package^58^ using weighted trimmed means normalization factors with condition (CNO vs. saline) as the main outcome of interest. Multiple testing correction was performed using the Benjamini-Hochberg approach to control for the false discovery rate (FDR)^59^. Gene ontology (GO) analyses were performed using Entrez gene IDs with the clusterProfiler R Bioconductor package^60^.

### RiboTag Experiments

For analysis of gene expression in vHC-PrL projectors, Rpl22^HA^/RiboTag animals were killed, brains were extracted, and the ventral portion of the HC was dissected out and flash frozen. Ribosome-mRNA complexes were affinity purified using a mouse monoclonal HA antibody (MMS-101R, Covance, Princeton, NJ) and A/G magnetic beads (88803 Pierce). RNA from IP samples was purified using RNeasy microcolumns (Qiagen, Valencia, CA) and quantified using the Ribogreen RNA assay kit (R11490, Invitrogen). Sequencing libraries were prepared using the Nugen Ovation SoLo RNAseq System and sequenced on a HiSeq2000 (Illumina). Differential gene expression and GO analyses were performed on IP samples from the projector group and the syn1-expressing vHC neuron group as described above.

### Statistical Analyses

Individual statistical tests are noted throughout figure legends in the text. All statistical tests, except where noted, were performed with GraphPad Prism software. Non-parametric tests were chosen when dependent variables could not be assumed to have a normal distribution. An alpha level of 0.05 was used to determine statistical significance.

## Acknowledgements

The authors wish to thank Richard de los Santos Abreu, Madhavi Tippani, Danisha Gallop, Leon Lin, and Martha Kimos for technical assistance. The authors additionally wish to thank Daniel Weinberger for invaluable comments on earlier versions of the manuscript. Funding for these studies was provided by the Lieber Institute for Brain Development, a National Institute of Mental Health R01 to KM (MH105592), and a National Institute of Mental Health F32 to HH (MH121052-01).

## Contributions

Study design: H.H., K.R.M. K.M. Behavioral experiments: H.H., H.Q., Y.M. Calcium imaging: H.H., H.Q. RiboTag experiments: H.H., K.R.M. RNA-sequencing: J.H.S. *Ex vivo* electrophysiology: G.H., H-Y.C., B.M. Data analysis: H.H., K.R.M., A.J. Writing leads: H.H., K.M.

