## Supplementary_Figures for "Molecularly-Defined Hippocampal Inputs Regulate Population Dynamics in the Prelimbic Cortex to Suppress Context Fear Memory Recall"

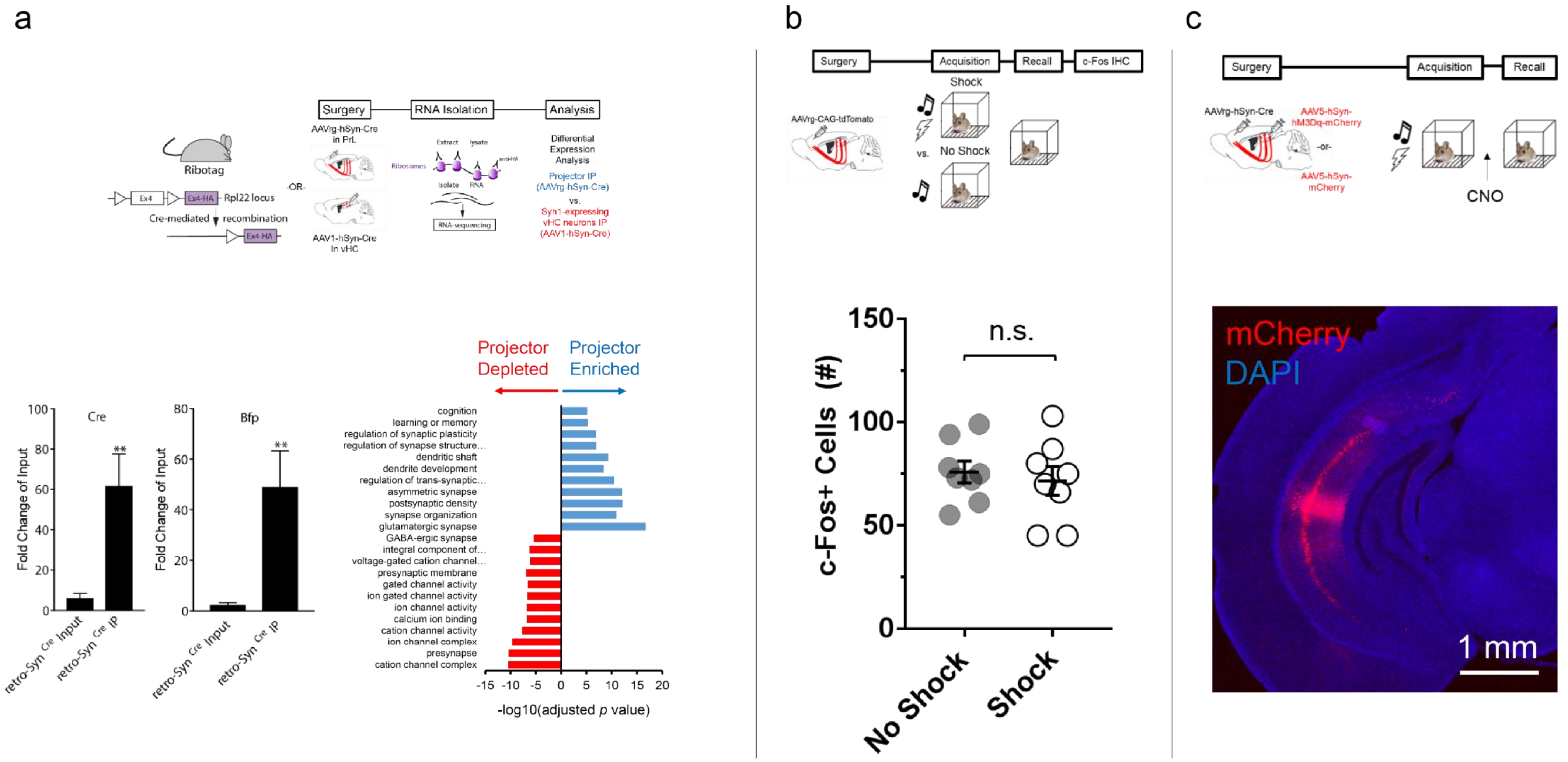


**Supplementary Figure 1.** *(a) Cre* ($t(10) = 3.509$, $p = 0.0056$) and *Bfp* ($t(10) = 3.219$, $p = 0.0092$; paired t-tests) are both enriched in IP samples from the vHC in Rpl22$^{HA}$/RiboTag mice in the projector group (left panel). Gene ontology (GO) terms for significantly enriched gene sets in the projector (blue bars) and syn1-expressing vHC neuron (red bars) groups (right panel). *(b)* Following context recall, total number of c-Fos+ neurons in the vHC does not differ between mice that received a shock during conditioning, and mice that did not receive a shock during conditioning ($t(14) = 0.5134$, $p = 0.6157$ for unpaired t-test). *(c)* Cre-dependent injections of either AAV5-hSyn-hM3Dq-mCherry or AAV5-hSyn-mCherry into the vHC result in mCherry expression that is largely limited to the CA1 subfield of the vHC.

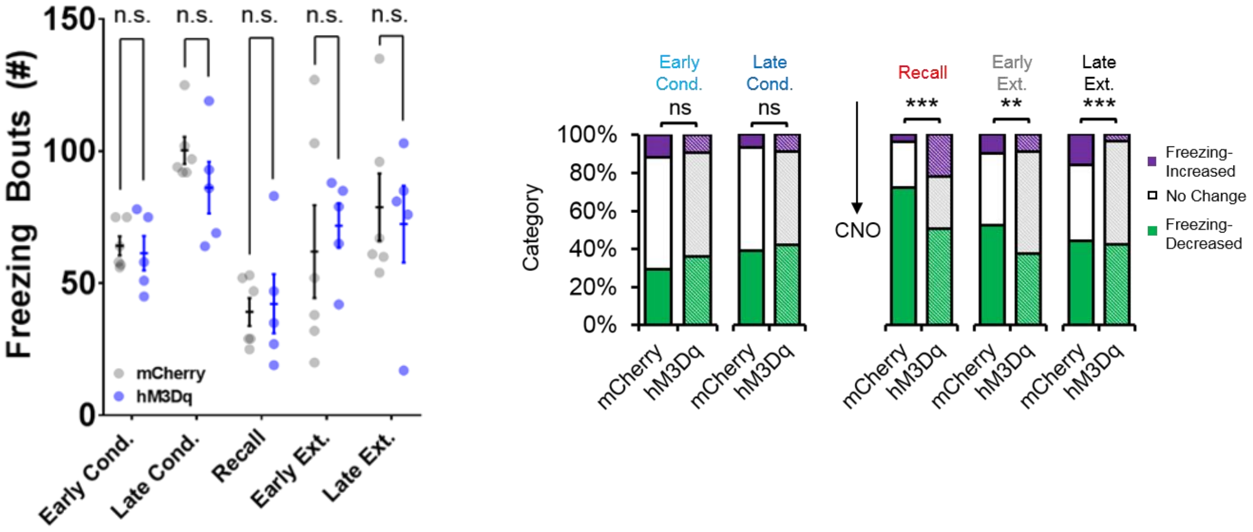


**Supplementary Figure 2.** *(a)* Unilateral activation of vHC-PrL projectors does not affect number of freezing bouts during any phase of fear conditioning ($F(4,36) = 0.38$, $p = 0.82$ for group by fear phase interaction, $F(1,9) = 0.098$, $p = 0.76$ for main effect of group, mixed-factorial ANOVA). *(b)* The proportion of PrL neurons that show either freezing-increased, freezing-neutral, or freezing-decreased responses does not differ between mCherry and hM3Dq groups during conditioning. These proportions, however, significantly differ between mCherry and hM3Dq groups during context recall ($p < 0.0001$), early extinction ($p = 0.004$), and late extinction ($p < 0.0001$; $X^2$ tests).

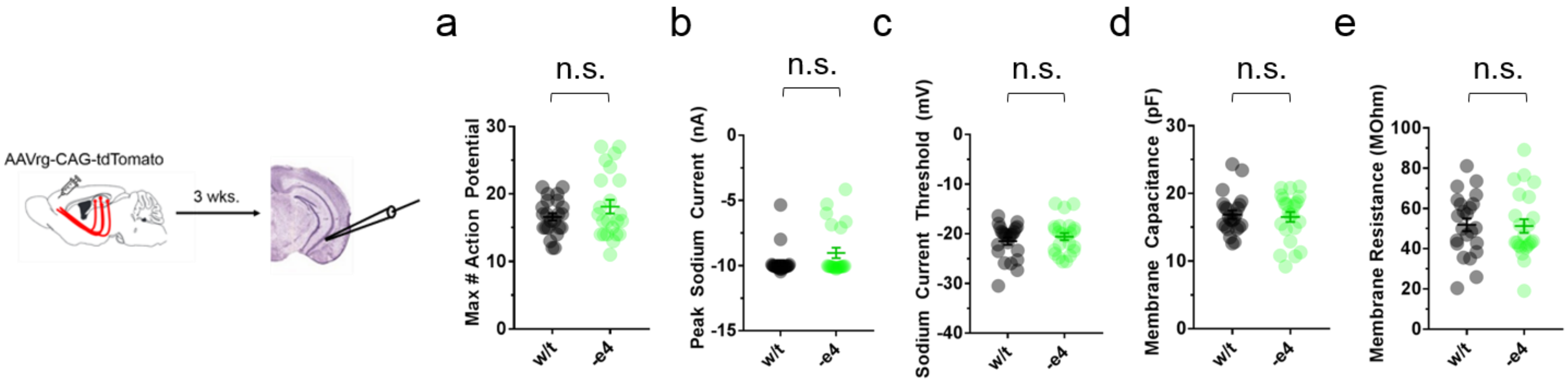


**Supplementary Figure 3.** *(a)* Maximum number of action potentials ($t(44) = 1.32$, $p = 0.196$), *(b)* peak sodium current ($t(44) = 1.7$, $p = 0.097$), *(c)* sodium current threshold ($t(44) = 0.88$, $p = 0.38$), *(d)* membrane capacitance ($t(44) = 0.36$, $p = 0.72$), and *(e)* membrane resistance ($t(44) = 0.14$, $p = 0.89$) in vHC-PrL projectors are not significantly different between w/t and –e4 mice (unpaired t-tests).

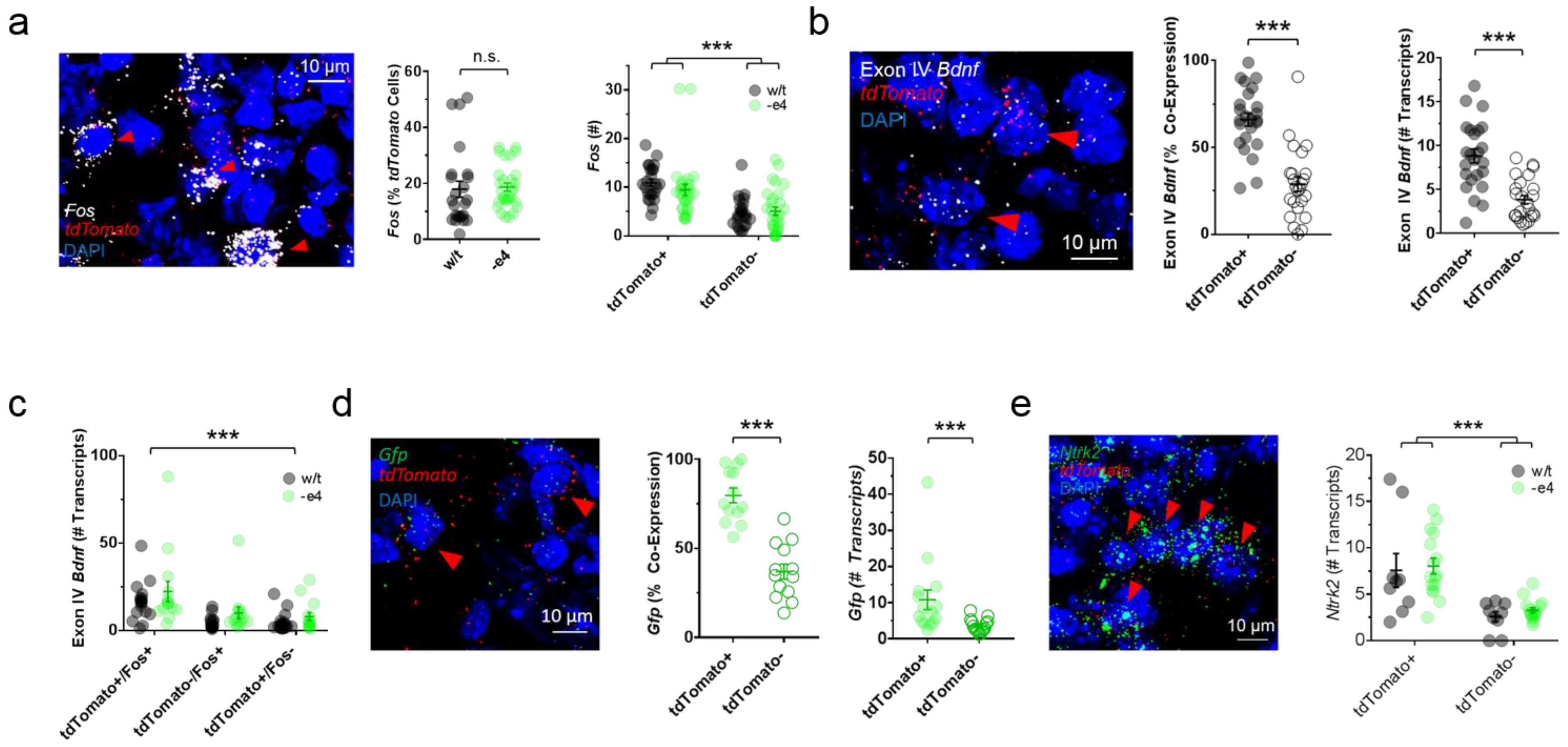


**Supplementary Figure 4.** *(a)* Neither percent co-expression ($t(51) = 0.25$, $p = 0.81$, unpaired t-test; left panel), nor number of *Fos* transcripts per neuron ($F(1,51) = 1.33$, $p = 0.25$ for genotype by cell type interaction; $F(1,51) = 46.37$, $p < 0.0001$, main effect of cell type, two-way ANOVA; right panel) in vHC-PrL projectors differs between w/t and –e4 mice following context fear recall. *(b)* vHC-PrL projectors co-express exon IV-containing *Bdnf* at significantly higher levels than non-projectors ($t(23) = 6$, $p < 0.0001$; left panel), and contain significantly more exon IV-containing *Bdnf* transcripts than non-projectors ($t(23) = 4.94$, $p < 0.0001$, paired t-tests; right panel). *(c)* vHC-PrL projectors recruited during context fear recall contain significantly more exon IV-containing *Bdnf* transcripts than recruited non-projectors and non-recruited projectors in both genotypes ($F(2,60) = 28.37$, $p < 0.0001$, main effect of cell type for mixed-factorial ANOVA). *(d)* vHC-PrL projectors co-express *Gfp* at significantly higher levels than non-projectors ($t(12) = 8$, $p < 0.0001$; left panel), and contain significantly more *Gfp* transcripts than non-projectors ($t(14) = 2.58$, $p = 0.02$, paired t-tests; right panel) in –e4 mice. *(e)* vHC-PrL projectors contain significantly more *Ntrk2* transcripts than non-projectors in both genotypes ($F(1,23) = 35.43$, $p < 0.0001$, main effect of cell type, two-way ANOVA).
